# Vac8-driven vacuolar remodeling primes mega-autophagy and safeguards gamete developmental competence

**DOI:** 10.64898/2026.10.05.756849

**Authors:** Tianyao Xiao, Benjamin S. Styler, Elçin Ünal

**Affiliations:** Department of Molecular and Cell Biology, University of California, Berkeley, CA 94720, USA; California Institute for Quantitative Biosciences (QB3), University of California, Berkeley, Berkeley, California 94720, USA; Center for Computational Biology, University of California, Berkeley, Berkeley, California 94720, USA

## Abstract

During gametogenesis, organelle remodeling is coordinated alongside chromosome segregation to ensure gamete quality. In *Saccharomyces cerevisiae*, the precursor-cell vacuole permeabilizes following completion of meiotic divisions, releasing hydrolytic enzymes that degrade cellular material excluded from gametes. The regulation and biological significance of this developmentally-programmed process, termed mega-autophagy, remain poorly understood. Here we identify the conserved armadillo repeat-containing protein Vac8 as a key regulator of mega-autophagy. The presence of Vac8 during meiosis is required for timely execution of mega-autophagy. Specifically, during anaphase II, precursor-cell vacuoles undergo Vac8-dependent fusion to form fewer, larger structures. Deleting *VAC8* disrupts this remodeling and impairs subsequent vacuolar permeabilization, establishing meiotic vacuolar fusion as a prerequisite for mega-autophagy. Defective mega-autophagy compromises the ability of gametes to exit dormancy and proliferate upon nutrient repletion. Together, these findings reveal that Vac8-mediated vacuolar remodeling during meiosis primes a post-meiotic quality-control mechanism that safeguards gamete developmental competence.

**Summary:** Xiao et al. show that Vac8 promotes vacuolar fusion during meiosis, priming the vacuole for post-meiotic lysis and clearance of cellular material excluded from gametes. Their findings link meiotic vacuolar remodeling to gamete developmental competence, with implications for germline quality control.

## Introduction

Sexual reproduction relies on the generation of healthy gametes. Gametes are produced through a specialized developmental program, termed gametogenesis, comprised of meiotic divisions followed by post-meiotic gamete development (Larose et al., 2019; Bergero et al., 2021). Quality control mechanisms during gametogenesis that partition essential cellular components into developing gametes while excluding unwanted materials are indispensable for safeguarding germline and offspring integrity, supporting production of a new generation despite age- or stress-associated defects in progenitors (Yanowitz, 2010; Sing et al., 2022; Ito and Shinohara, 2022; Xiao and Ünal, 2025). While most studies of meiosis have focused on cell-cycle regulation and chromosome remodeling, defining how organelles and cytoplasmic contents are remodeled during gametogenesis is essential for understanding the cellular basis of gamete quality control (Jones, 2007; Yamashita, 2023).

Studies in the budding yeast *Saccharomyces cerevisiae* have provided direct evidence for a resetting of gamete homeostasis through meiotic differentiation. Regardless of the progenitor cell’s replicative age, gametes–also known as spores–are born young and free of parental age-associated factors (Unal et al., 2011; King et al., 2019; Schneider et al., 2024; Spiri et al., 2025).

For instance, nuclear-localized senescence-associated factors are separated from gametes through a five-way nuclear division during meiosis II. While sister chromatids are segregated to form four gamete nuclei, nuclear senescence-associated factors and the core subunits of the nuclear pore complex (NPC) are sequestered into a membrane-bound compartment that is ultimately excluded from gametes. This structure is called the Gametogenesis Uninherited Nuclear Compartment (GUNC) (King et al., 2019). Concurrently, gamete plasma membrane is synthesized de novo within the precursor cell, initiating from the spindle pole body near each gamete nuclei (Moens, 1971; Neiman, 1998, 2024). This developing membrane sac, termed the prospore membrane (PSM), packages adjacent gamete nuclei and restricts GUNC substrates from entering into gametes (King et al., 2019). At the same time, mitochondria and the endoplasmic reticulum (ER) also detach from the cell cortex and are enriched around the GUNC, forming a super-organellar structure (Miyakawa et al., 1984; Suda et al., 2007; King et al., 2019; Sawyer et al., 2019; Otto et al., 2021; Ruediger et al., 2025). Interestingly, NPCs undergo similar remodeling during metazoan gametogenesis (Ho, 2010; Li et al., 2023; Thomas et al., 2023; Santos et al., 2024; Venkat et al., 2026), and compartmentalization and restricted inheritance of mitochondria also occur in oogenesis and spermatogenesis (DeLuca and O’Farrell, 2012; Lieber et al., 2019; Varuzhanyan et al., 2019; Palozzi et al., 2022; Schwartz et al., 2022; Ayachit and Shravage, 2023; Zhang et al., 2024). These features underscore the broad conservation of organelle quality control mechanisms during gametogenesis.

At the end of meiosis, the four PSMs close through cytokinesis, leaving the GUNC and other uninherited organellar and cytosolic contents in the anucleate precursor cell, termed the ascus (Neiman, 1998, 2011; King et al., 2019). The precursor-cell vacuoles are fully excluded from the gametes which instead synthesize their vacuoles de novo post-meiotically (Roeder and Shaw, 1996; Eastwood and Meneghini, 2015). Once gametes are fully mature with a protective spore wall, the precursor-cell vacuole ruptures, releasing hydrolytic enzymes to degrade leftover cellular contents (Eastwood et al., 2012; Eastwood and Meneghini, 2015; King et al., 2019). This event is termed mega-autophagy (Eastwood et al., 2012). Intriguingly, analogous clearance processes have been described in nematode and mouse oocytes (Bohnert and Kenyon, 2017; Samaddar et al., 2021; Zaffagnini et al., 2024; Satouh et al., 2025) and metazoan spermatogenesis (Breucker et al., 1985; Blanco-Rodríguez and Martínez-García, 1999; Huang et al., 2012; Lv et al., 2020; Lei et al., 2021; Arslan et al., 2026). However, the regulatory mechanisms coordinating this post-meiotic degradation process, and its biological significance remain unclear.

From a genetic screen, we identified *VAC8* as a factor required for mega-autophagy. We further characterized how Vac8 regulates mega-autophagy and found that Vac8-mediated vacuolar fusion during late meiosis serves as a priming step for the post-meiotic lysis. Using mutants lacking Vac8, we were also able to explore how Vac8 and mega-autophagy affects spores, especially their ability to exit dormancy and re-enter mitotic growth in response to nutrient signals. Our results highlight a significant regulatory role of Vac8 in vacuolar remodeling during meiosis and link meiotic vacuolar quality control with gamete developmental competence.

## Results

### Vac8 is required for the post-meiotic degradation of uninherited contents

We previously established an imaging-based genetic screening strategy using a nuclear-localized genetically encoded multimeric protein (nucGEM) as a reporter for GUNC-mediated exclusion and clearance (Szoradi et al., 2022; Hernandez et al., 2024; Ruediger et al., 2025). nucGEM forms discrete nuclear agglomerates that behave similarly to endogenous GUNC substrates: they are sequestered away from gamete nuclei during anaphase II, separated from the developing gametes by cytokinesis, and subsequently degraded by mega-autophagy (Ruediger et al., 2025). Here, we used this reporter to perform an imaging-based screen of the SK1 non-essential gene deletion collection to identify factors required for either gamete exclusion or post-meiotic clearance of uninherited cellular material. The SK1 background is highly competent for sporulation in that the vast majority of cells complete meiosis and spore maturation within 24 h upon transfer to sporulation medium (SPO) (Fig. 1A) (Kane and Roth, 1974). We therefore acquired endpoint images and quantified the fraction of sporulated cells that retain nucGEM foci. Remarkably, this genome-wide screen identified a single mutant, *vac8*Δ, in which nucGEMs were efficiently excluded from gametes but failed to undergo subsequent degradation, pointing to a specific requirement for Vac8 in mega-autophagy.

**Figure 1.**
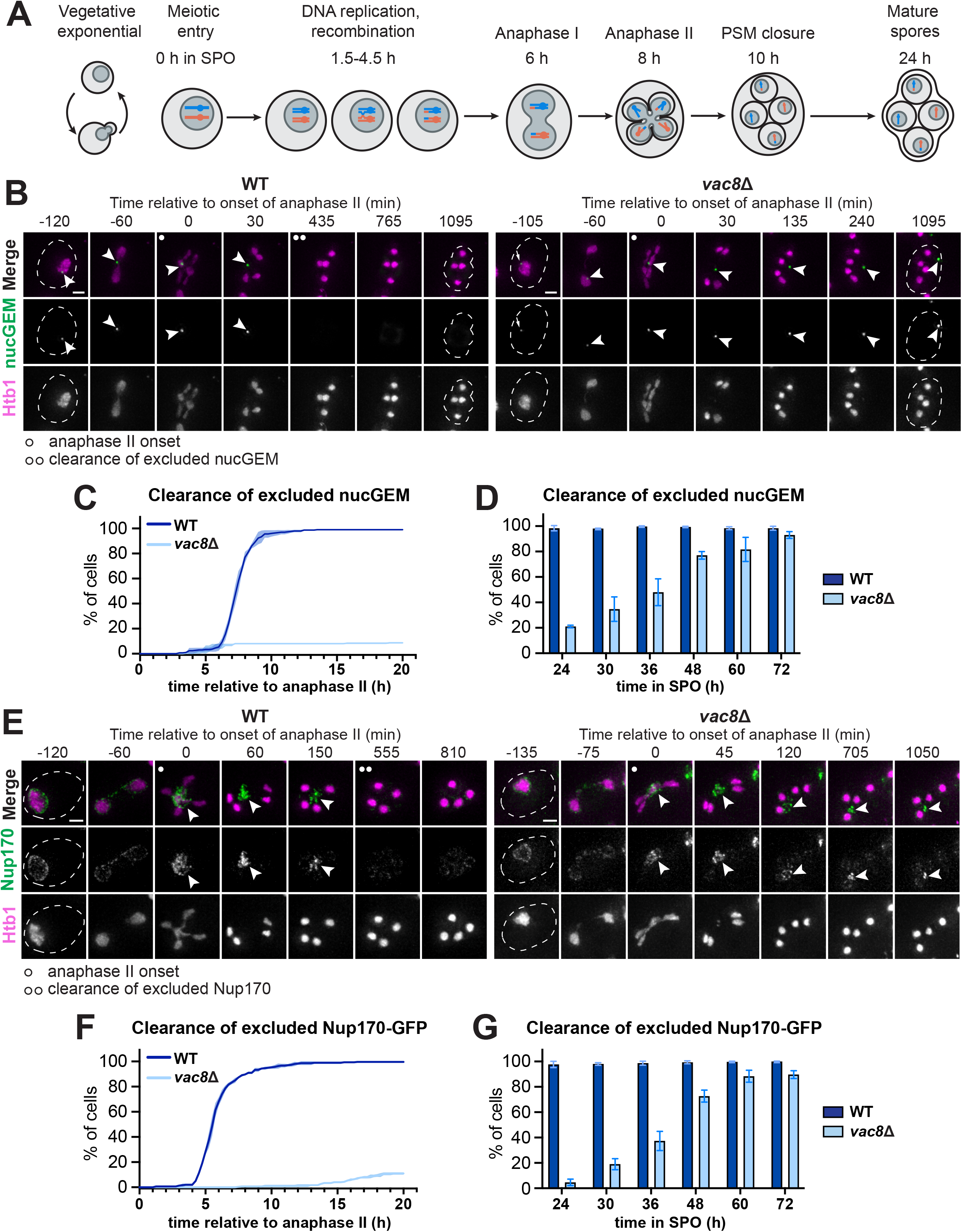
Vac8 is required for post-meiotic degradation of uninherited GUNC substrates. **(A)** Schematic of the meiotic timeline in the SK1 strain background. When nutrients are adequate, diploid budding yeast undergoes vegetative growth through mitotic division. When switched to sporulation medium (SPO) in which the carbon source is acetate, cells enter meiosis. SK1 strains enter anaphase I after around 6 h in SPO and anaphase II after around 8 h in SPO. Spores are packaged at about 10 h in SPO and continue to develop until maturation. **(B)** Montage of wild-type (WT) or *vac8*Δ cells expressing nucGEM (white arrowheads) driven by the *ATG8* promoter progressing through gametogenesis. Chromatin was visualized with the histone marker Htb1-mCherry. The time point depicting anaphase II onset was defined as 0 min and indicated by the single white dot on the top left corner of the merged image. The time point depicting nucGEM clearance is indicated by the double white dot on the top left corner of the merged image. Images are shown as maximum-intensity projections (MIPs) of z-stacks. Scale bar = 2 µm. **(C)** Quantification of (B) showing the percentage of WT and *vac8*Δ cells that had cleared uninherited nucGEM plotted relative to the time of anaphase II onset for each cell. Shaded regions indicates SD of three replicates, n = 100 cells per replicate. **(D)** The percentage of WT and *vac8*Δ cells that had cleared uninherited nucGEM at indicated time points in SPO. Error bars = SD of three replicates, n = 100 cells per replicate. **(E)** Montage of WT and *vac8*Δ cells expressing endogenously labelled Nup170-GFP progressing through gametogenesis. Chromatin was visualized with Htb1-mCherry. The time point depicting anaphase II onset was defined as 0 min and indicated by the single white dot on the top left corner of the merged image. White arrowheads indicate Nup170-GFP excluded from gametes. The time point depicting the clearance of excluded Nup170-GFP was indicated by the double white dot on the top left corner of the merged image. Images are shown as MIPs of z-stacks. Scale bar = 2 µm. **(F)** Quantification of (E) showing the percentage of WT and *vac8*Δ cells that had cleared excluded Nup170-GFP plotted relative to the time of anaphase II onset for each cell. Shaded region indicates SD of three replicates, n = 100 cells per replicate. **(G)** The percentage of WT and *vac8*Δ cells that had cleared excluded Nup170-GFP at indicated time points in SPO. Error bars = SD of three replicates, n = 100 cells per replicate.

*VAC8* encodes the armadillo-repeat containing (ARM) protein Vac8, which localizes to the vacuolar membrane upon lipid modification (Subramanian et al., 2006; Nadolski and Linder, 2009; Lei et al., 2021; Kim et al., 2023; Popelka and Klionsky, 2025). Vac8 acts as an inter-organelle adaptor for vacuoles in mitotic cells, mediating the interaction between vacuoles and other compartments, including the nucleus and ER (Pan et al., 2000; Murley et al., 2015; Jeong et al., 2017), lipid droplets (Diep et al., 2024; Álvarez-Guerra et al., 2024), myosin (Tang et al., 2003; Wong et al., 2020), and autophagosome machinery (Hollenstein et al., 2019; Gatica et al., 2021; Lei et al., 2021; Hollenstein et al., 2021). The deletion of *VAC8* abolishes sporulation in S288C-derived strains, potentially due to defects in autophagy that impair meiotic entry (Enyenihi and Saunders, 2003; Barve and Manjithaya, 2021). Because of this, the precise roles of Vac8 during meiosis have not been previously explored. Notably, Vac8 is conserved among metazoans, and its orthologs are extensively expressed in germline tissues (Fouchécourt et al., 2019; Lei et al., 2021). In addition, mutations in Vac8 orthologs are linked to impaired gamete function and infertility in multiple organisms (Cheng et al., 2013; Pausch et al., 2016; Lei et al., 2021; Rahim et al., 2024). Unlike the well-defined vacuolar membrane localization of Vac8 in yeast, the cellular localization of its metazoan orthologs remains unclear. Nevertheless, these findings suggest a conserved role for Vac8 in germline quality control.

In the SK1 background, *VAC8* deletion did not impair meiotic entry or progression. Instead, *vac8*Δ cells progressed slightly faster through meiosis, initiating anaphase II 0.84 ± 0.16 h earlier than wild-type (WT) cells (Fig. S1, A and B). Overall sporulation efficiency was unaffected (94.0% for WT and 96.3% for *vac8*Δ cells), although the distribution of tetrads/triads and dyads/monads formed was slightly altered (Fig. S1C). This phenotype contrasts sharply with the S288C-derived BY background, in which *VAC8* deletion prevents meiotic entry (Enyenihi and Saunders, 2003; Barve and Manjithaya, 2021). This deficiency cannot be fully rescued by introducing known SK1-associated alleles that enhance sporulation efficiency (Harvey et al., 2018) into a “fixed” BY background (Fig. S1D), suggesting that additional genetic factors underlie the ability of SK1 *vac8*Δ cells to undergo meiosis. Importantly, the robust sporulation of SK1 *vac8*Δ cells enabled us to investigate the role of Vac8 during gametogenesis other than meiotic entry.

We first validated the screen result by confirming that *VAC8* deletion severely impairs the post-meiotic clearance of nucGEMs (Fig. 1, B-D), without affecting their exclusion from gametes (Fig. S1E). We also assessed nucGEM degradation by monitoring the appearance of a free GFP band (∼25kD) through immunoblotting, which provides a biochemical readout of substrate degradation (King et al., 2019). Consistent with the microscopy data, *VAC8* deletion markedly delayed post-meiotic nucGEM degradation (Fig. S1F). This defect was not restricted to the synthetic nucGEM substrate: degradation of the fluorescently tagged Nup170, an endogenous GUNC substrate (King et al., 2019), was similarly impaired (Fig. 1, E-G; Fig. S1, E and G). In addition to nuclear substrates, *VAC8* deletion also disrupted the post-meiotic clearance of proteins associated with other cellular compartments that are excluded from gametes, including Cit1 (mitochondria) (Sawyer et al., 2019), Sec63 (ER) (Otto et al., 2021), and Pma1 (plasma membrane) (Eastwood and Meneghini, 2015) (Fig. S1, H-I). Together, these findings identify Vac8 as a factor required for timely post-meiotic degradation of diverse cellular contents excluded from gametes.

### Vac8 is required for timely mega-autophagy and the developmental competence of resulting gametes

We next sought to determine whether *vac8*Δ disrupts clearance through a defect in mega-autophagy. Mega-autophagy is accomplished by the rupture and lysis of precursor-cell vacuoles following meiosis (Eastwood and Meneghini, 2015; Eastwood et al., 2012). To determine whether *VAC8* deletion affects vacuolar integrity, we visualized the precursor-cell vacuoles via FM4-64 staining prior to meiotic entry. FM4-64 is internalized through endocytosis and ultimately labels the vacuolar membrane (Vida and Emr, 1995). Because vacuoles in the precursor cell are not inherited by gametes, this approach specifically labels the precursor vacuolar compartment (Roeder and Shaw, 1996). After 24 h in SPO, over 95% of sporulated WT cells had completed mega-autophagy, and FM4-64 was redistributed from the lysed precursor vacuoles to the ascus space and/or spore walls (Roeder and Shaw, 1996), likely reflecting release of the dye following vacuolar membrane rupture (Fig. 2 A and B). In contrast, FM4-64 remained associated with the intact precursor vacuoles of *vac8*Δ cells (Fig. 2, A and B), indicating a defective post-meiotic vacuolar lysis that is consistent with the impaired clearance of uninherited cellular contents (Fig 1 B-G; Fig S1 F-I).

**Figure 2.**
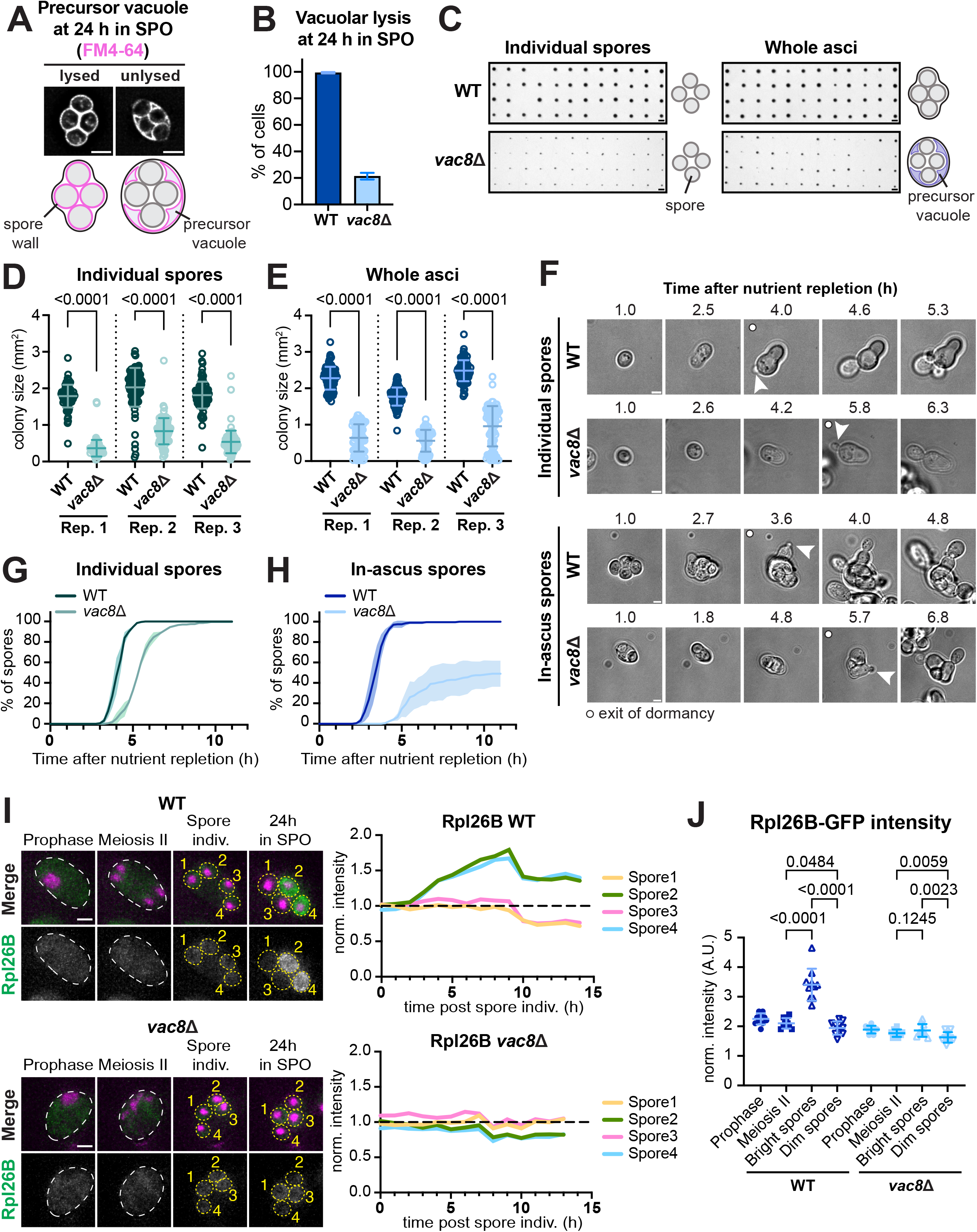
Vac8 is required for post-meiotic vacuolar lysis and spore development. **(A)** Widefield microscopy images of sporulated cells at 24 h in SPO. Vacuoles of the meiotic precursor cells were visualized by FM4-64 staining prior to switching to SPO. Unlysed precursor vacuoles retain FM-64. In asci with completed mega-autophagy, FM4-64 non-specifically stains the asci and spore wall. Image represents a single z-slice. Scale bar = 2 µm. In the schematic, FM4-64 staining is indicated in magenta. **(B)** The percentage of WT or *vac8*Δ cells that had lysed the precursor cell vacuole at 24 h in SPO. Error bars = SD of three replicates, n = 100 cells per replicate. **(C)** Colonies grown from individual spores or whole asci derived from sporulated WT or *vac8*Δ cells after being placed on nutrient-enriched (YPD) plates for two days. After 24 h in SPO, spores or whole asci were transferred to YPD plates. Scale bar = 2 mm. In the schematic, the precursor vacuole is shown in purple. **(D and E)** Quantification of (C) showing the size of colonies (mm^2^) derived from individual spores (D) or whole asci (E) on YPD plates. Each symbol represents the size of an individual colony, n > 80 colonies for each group. Error bars = SD of all individual colonies of the same genotype. Three independent pairs of experiments were compared and separated by dashed vertical lines. **(F)** Brightfield montage of free or in-ascus spores derived from sporulated WT or *vac8*Δ cell progressing through germination in nutrient-enriched liquid media (YPD). After 24 h in SPO, spores or whole asci were transferred to 96-well imaging plates and switched to YPD. The time point when switching to YPD was defined as 0 h. Germination was defined by the appearance of the first bud from each free spore or any spore in an entire ascus, as indicated by white arrowheads. The time point depicting germination was indicated by the white circle on the top left corner of the merged image. Image represents a single z-slice. Scale bar = 2 µm. **(G and H)** Quantification of (F) showing the percentage of free (G) or in-ascus (H) spores that had produced the first bud at each time point after switching to YPD. Shaded region indicates SD of three replicates, n = 100 spores per replicate. **(I)** Montage of WT and *vac8*Δ cells expressing endogenously labelled Rpl26B-GFP at indicated stages during gametogenesis. Chromatin was visualized with Htb1-mCherry (shown in magenta in merged images). The precursor cells or asci are outlined in white, and spores are outlined in yellow. Images are shown as MIPs of z-stacks. Scale bar = 2 µm. The line plots on the right depict normalized Rpl26B fluorescence intensity in individual spores at each time point after spore individualization. Each line represents the respective spore labelled in the microscopy image. The time point of spore individualization was defined as 0 h. Fluorescence intensity was normalized to that at 0 h, and the dashed black line indicates normalized intensity = 1.0. **(J)** Quantification of the normalized Rpl26B fluorescence intensity in WT or *vac8*Δ cells during prophase, anaphase II, and in bright versus dim spores after 24 h in SPO. Each symbol represents an individual cell or spore, n = 10 sporulated cells. Error bars = SD across all individual cells or spores within each group. For statistical analysis information related to (D), (E), and (J), see Table S5. The p-values are indicated in the graph.

It has been previously shown that defects in spore wall formation can inhibit mega-autophagy, presumably preventing premature exposure of unprotected gametes to vacuolar hydrolases (Eastwood and Meneghini, 2015). Therefore, we asked whether the mega-autophagy defect in *vac8*Δ cells could result from impaired spore wall development. We assessed the formation of the outermost dityrosine layer, which marks the completion of spore wall, by measuring its autofluorescence following NH₄OH alkalinization (Kungl et al., 1994; Suda et al., 2009a). WT spores displayed peripheral dityrosine autofluorescence, whereas control mutants lacking the dityrosine biosynthetic enzymes Dit1 and Dit2 (Briza et al., 1994) showed no detectable signal (Fig. S2, A and B). Similar to WT, *vac8*Δ spores also exhibited dityrosine autofluorescence, albeit at higher levels (Fig. S2, A and B). Thus, the failure of *vac8*Δ cells to execute mega-autophagy is unlikely a result of incomplete spore wall formation.

Given that mega-autophagy is a programmed event following meiotic division, we next asked whether loss of Vac8 affects the developmental competence of the resulting gametes. Nutrient deprivation induces diploid yeast cells to undergo meiosis and form dormant gametes, known as spores, which can re-enter mitotic growth upon nutrient replenishment (Joseph-Strauss et al., 2007). To assess their ability to resume growth, we first released individual spores from the ascus by zymolyase digestion and plated them on nutrient-rich medium. After two days, colonies derived from single *vac8*Δ spores were significantly smaller than those derived from WT spores (Fig. 2, C and D). When plated without zymolyase treatment, we observed an even stronger phenotype (Fig. 2, C and E). This latter condition more closely approximates physiological germination, in which spores initiate growth in response to nutrient availability and subsequently emerge through the ascus wall, without prior artificial enzymatic removal of the surrounding material.

To determine whether the reduced colony growth reflected a delayed exit from dormancy, or a decrease in subsequent growth, we monitored the first budding event following nutrient replenishment by time-lapse imaging (Hugener et al., 2024; Ruediger et al., 2025; Reyes et al., 2025). Individual *vac8*Δ spores initiated budding approximately 2 h later than WT spores (Fig. 2, F and G; Fig. S2C), representing a substantial delay. The defect was more pronounced for spores retained within intact asci, with approximately 40% of *vac8*Δ asci failing to initiate budding during the entire imaging period (Fig. 2, F and H; Fig. S2D). Together, these results indicate that Vac8 is required for efficient exit from spore dormancy and re-entry into mitotic growth.

We next examined spore-autonomous protein expression as an additional measure of spore maturation. It has been shown that newly formed spores autonomously synthesize ribosomal proteins (Eisenberg et al., 2018). To monitor this process, we used heterozygous diploid cells carrying a single fluorescently-tagged copy of either *RPL26B* or *RPL29.* Spore-autonomous expression can be detected by the resulting 2:2 pattern of bright and dim spores within individual asci (Eisenberg et al., 2018). In *vac8*Δ cells, the spore-autonomous expression of both ribosomal protein reporters was markedly impaired relative to WT (Fig. 2, I and J; Fig. S2, E and F). Altogether, these findings establish that Vac8 is required not only for post-meiotic mega-autophagy but also for the developmental competence of the resulting gametes.

### Vac8 acts during meiosis to enable subsequent mega-autophagy

To explore how Vac8 functions to promote mega-autophagy, we first examined the expression and localization of GFP-tagged Vac8 during gametogenesis. Vac8 localized to precursor-cell vacuolar membranes, and its abundance increased as cells progressed through meiosis (Fig. 3A; Fig. S2G). Following spore packaging, Vac8 was also readily detected within newly generated spores (Fig. 3A; Fig. S2G). To distinguish whether Vac8 is required for mega-autophagy in precursor cells or newly formed spores, we used an auxin-inducible degron system to conditionally deplete Vac8 at defined stages of gametogenesis (Fig. 3B) (Nishimura et al., 2009;

**Figure 3.**
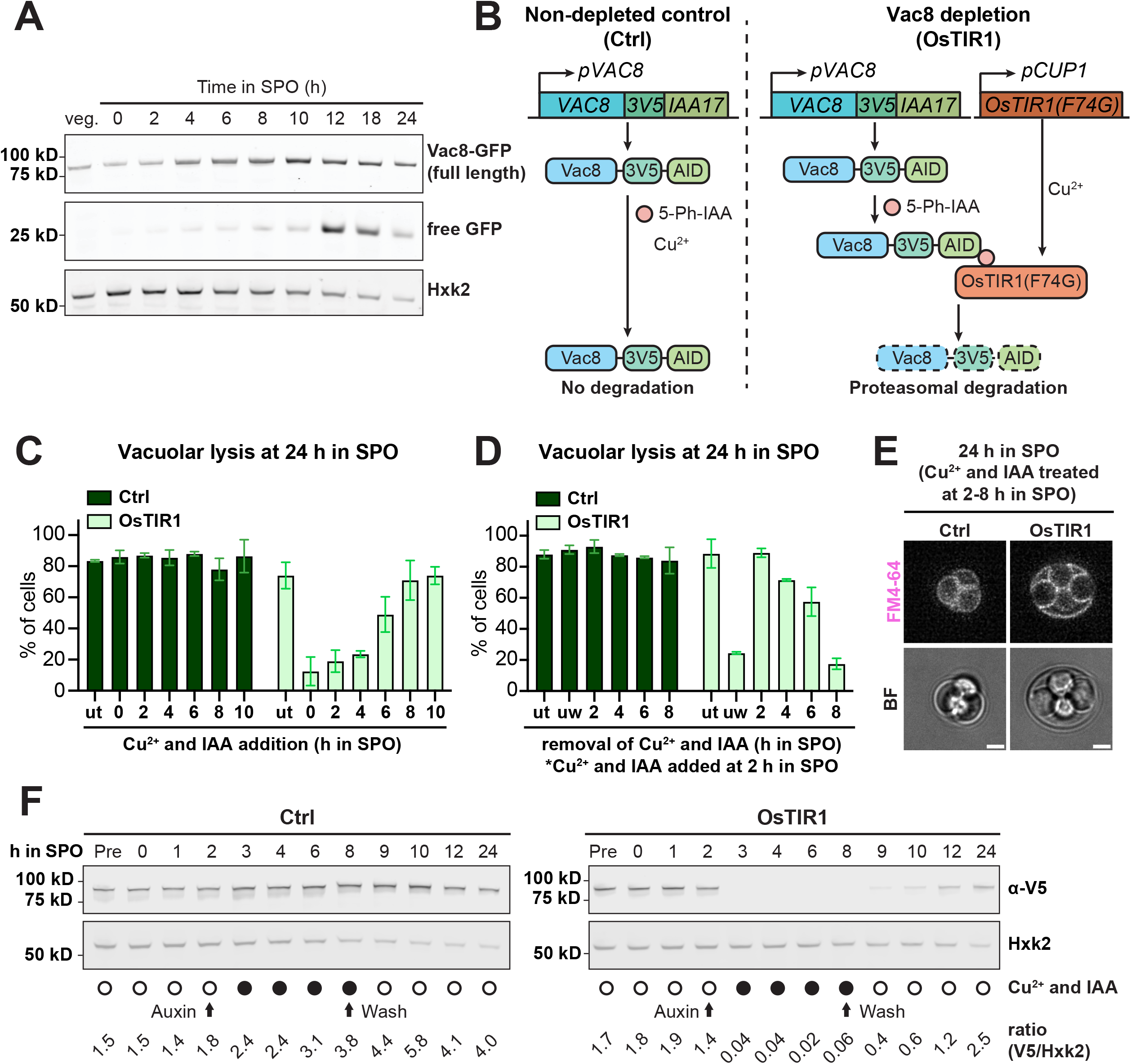
The presence of Vac8 during meiosis is critical for mega-autophagy. **(A)** Western blot analysis of Vac8-GFP (full length and free GFP) in WT cells undergoing vegetative growth (veg.) or progressing through gametogenesis. Samples were collected at indicated time points in SPO. Hexokinase 2 (Hxk2) was used as a loading control. **(B)** Schematic of the auxin-degron system that was used for Vac8 depletion during gametogenesis. Vac8 was endogenously tagged with 3V5-IAA17. In Vac8-depleted strains (OsTIR1), *OsTIR1(F64G)* gene was incorporated into the genome and driven by the Cu^2+^-inducible promoter *CUP1*. With Cu^2+^ (CuSO_4_) and IAA (5-Ph-IAA) addition, OsTIR1(F64G) interacts with the AID tag, mediating the proteasomal degradation of Vac8. The non-depleted control (Ctrl) lacks *OsTIR1(F64G)* gene allele, and thus the addition of Cu^2+^ and IAA could not cause Vac8-AID degradation. **(C)** The percentage of Ctrl or OsTIR1 cells that had lysed their precursor vacuoles after 24 h in SPO. Cu^2+^ and IAA were added to the sporulating culture at indicated time points in SPO. ut = untreated. Error bars = SD of two replicates, n = 100 cells per replicate. **(D)** The percentage of Ctrl or OsTIR1 cells that had lysed their precursor vacuoles after 24 h in SPO. Cu^2+^ and IAA was added to the sporulating culture at 2 h in SPO. At indicated time points, cells were washed and resuspended in conditioned SPO. Ut = untreated; uw = unwashed. Error bars = SD of two replicates, n = 100 cells per replicate. **(E)** Widefield microscopy images of sporulated Ctrl or OsTIR1 cells at 24 h in SPO. Meiotic precursor cell vacuoles were visualized by FM4-64 staining prior to switching to SPO. Cu^2+^ and IAA were added to the sporulating culture at 2 h in SPO, and cells were washed and resuspended in conditioned SPO at 8 h in SPO. Image represents a single z-slice. BF, bright field. Scale bar = 2 µm. **(F)** Western blot analysis of Vac8-3V5 in Ctrl or OsTIR1 cells progressing through gametogenesis. Samples were collected at indicated time points in SPO. Cu^2+^ and IAA were added to sporulating culture at 2h in SPO, and cells were washed and resuspended in conditional SPO at 8 h in SPO. Hxk2 was used as a loading control. Pre, pre-meiotic culture. The ratio of V5 to Hxk2 band intensity in each sample is indicated at the bottom.

Yesbolatova et al., 2020; Ruediger et al., 2025). Upon auxin addition, Vac8 abundance is reduced by >95% within 1.5 h during meiosis (Fig. S2H).

We next depleted Vac8 at different stages of gametogenesis and assessed post-meiotic precursor-vacuole lysis by FM4-64 staining (Fig. 2A). Depletion during early meiosis (prior to 4 h in SPO) recapitulated the mega-autophagy defect observed in in *vac8*Δ cells (Fig. 2B; Fig. 3C). In contrast, depletion after cells had entered meiosis II (8 h in SPO) had little effect on mega-autophagy relative to non-depleted controls (Fig. 3C). These results suggest that Vac8 executes its function in mega-autophagy before or during meiosis II.

To further define this temporal requirement, we depleted Vac8 beginning at 2 h in SPO and subsequently removed the depletion agents at different stages to restore Vac8 expression. Restoring Vac8 prior to anaphase II (8 h in SPO) rescued post-meiotic vacuolar lysis, whereas restoring Vac8 after anaphase II onset failed to rescue the mega-autophagy defect (Fig. 3 D and E). Immunoblotting confirmed that when Vac8 was depleted from 2-8 h in SPO, it was nearly undetectable during meiosis but reaccumulated after spore packaging (Fig. 3F). Despite this post-meiotic re-expression, these cells remained defective in mega-autophagy (Fig. 3 D-F). Together, these findings indicate that Vac8 functions in the precursor cell during meiosis, before anaphase II, to enable the subsequent post-meiotic lysis of vacuoles and trigger mega-autophagy.

### Mega-autophagy promotes efficient exit from spore dormancy

Our finding that Vac8 acts during meiosis to enable subsequent mega-autophagy provided an opportunity to distinguish the consequences of defective mega-autophagy from those caused by the continual absence of Vac8. We therefore transiently depleted Vac8 at 2-8 h in SPO to assess how failure to timely execute mega-autophagy affects the developmental competence of the resulting spores. Intriguingly, when spores were enzymatically released from asci and plated individually on nutrient-rich medium, spores derived from Vac8-depleted and control cells formed colonies of comparable size (Fig. 4 A and B). In contrast, when intact asci were plated without enzymatic disruption, the asci that had failed to undergo mega-autophagy produced significantly smaller colonies than non-depleted controls (Fig. 4 A and C). We next monitored dormancy exit directly by time-lapse microscopy following nutrient replenishment. Consistent with the colony-growth assays, individual spores released from asci exited dormancy with similar kinetics regardless of Vac8 depletion during meiosis (Fig. 4 D, E, and G). Strikingly, spores retained within asci that had failed to undergo mega-autophagy exhibited a pronounced defect in dormancy exit (Fig. 4 D, F, and H). These findings indicate that post-meiotic clearance of uninherited cellular material is important for efficient germination when spores remain within their native environment.

**Figure 4.**
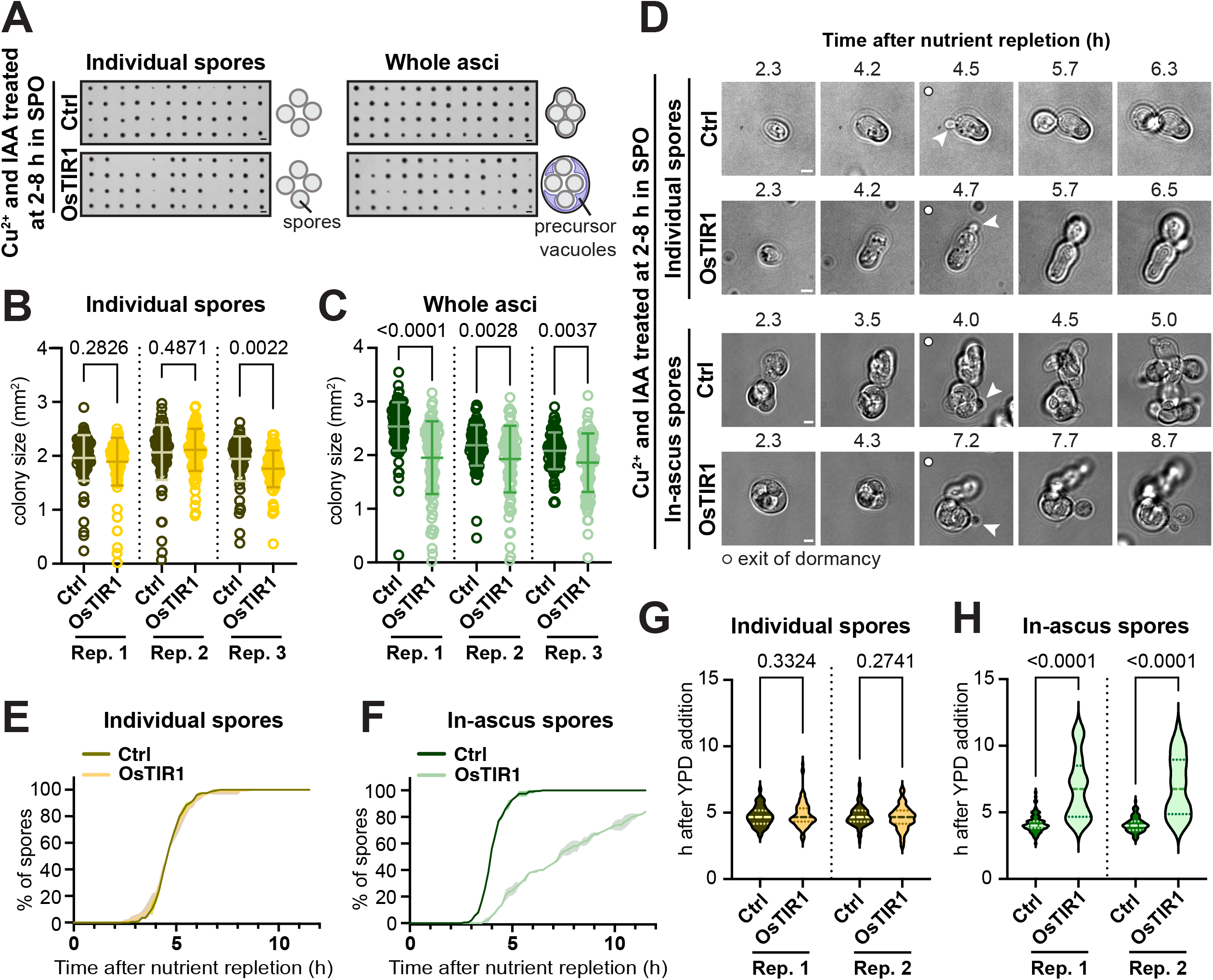
Mega-autophagy is a prerequisite for in-ascus spores to exit dormancy upon nutrient repletion. **(A)** Colonies grown from individual spores or whole asci derived from sporulated Ctrl or OsTIR1 cells after being placed on YPD plates for two days. During sporulation, Cu^2+^ and IAA were added to the culture at 2h in SPO, and cells were washed and resuspended in conditional SPO at 8 h in SPO. After 24 h in SPO, spores or whole asci were transferred to YPD plates. Scale bar = 2 mm. In the schematic, the precursor-cell vacuole is shown in purple. **(B and C)** Quantification of (A) showing the size of colonies (mm^2^) derived from individual spores (B) or whole asci (C) on YPD plates. Each symbol represents the size of an individual colony, n > 80 colonies for each group. Error bars = SD of all individual colonies of the same genotype. Three independent pairs of experiments were compared and separated by dashed vertical lines. **(D)** Brightfield montage of free or in-ascus spores derived from sporulated Ctrl or OsTIR1 cells progressing through germination in YPD. During sporulation, Cu^2+^ and IAA were added to the culture at 2h in SPO, and cells were washed and resuspended in conditional SPO at 8 h in SPO. After 24 h in SPO, spores or whole asci were transferred to 96-well imaging plates and switched to YPD. The time point when switching to YPD was defined as 0 h. Germination was defined by the appearance of the first bud from each free spore or any spore in an entire ascus, as indicated by white arrowheads. The time point depicting germination was indicated by the white circle on the top left corner of the merged image. Image represents a single z-slice. Scale bar = 2 µm. **(E and F)** Quantification of (D) showing the percentage of free (E) or in-ascus (F) spores that had produced the first bud at each time point after switching to YPD. Shaded region indicates SD of three replicates, n = 100 spores per replicate. **(G)** Quantification of (D) showing the time it took for free or in-ascus spores to germinate after switching to YPD. n = the number of spores that had germinated in (E) and (F). For statistical analysis information related to (B), (C), (G), and (H), see Table S5. The p-values are indicated in the graph.

The phenotype caused by meiosis-specific Vac8 depletion was less severe than that of *vac8*Δ cells, despite similar defects in mega-autophagy. Whereas individual *vac8*Δ spores exhibited delayed dormancy exit and reduced colony growth (Fig. 2, C-H), spores in which Vac8 was restored following meiosis behaved similarly to controls once enzymatically released from their enclosing structure (Fig. 4, A-F). This distinction suggests that Vac8 expressed in newly formed spores contributes to spore fitness, whereas its meiotic function promotes mega-autophagy and efficient dormancy exit under more physiological conditions. Together, these results indicate that developmental clearance of uninherited cellular contents facilitates the ability of spores to resume mitotic growth in response to nutrient repletion within their native environment.

### Vac8 promotes mega-autophagy independently of its known tethering functions

To investigate how Vac8 promotes mega-autophagy, we first asked whether this function could be explained by its previously characterized roles. In mitotically dividing cells, Vac8 functions as a vacuolar membrane-associated adaptor that establishes interactions between vacuoles and other cellular structures (Fig. 5A) (Subramanian et al., 2006; Lei et al., 2021; Kim et al., 2023; Álvarez-Guerra et al., 2024; Popelka and Klionsky, 2025). Through its interaction with Vac17, Vac8 links vacuoles to myosin and mediates vacuolar inheritance during mitosis (Wang et al., 1998; Pan and Goldfarb, 1998; Tang et al., 2003; Wong et al., 2020; Kim et al., 2023). Vac8 also forms inter-organelle contacts, including the nucleus-vacuole junction (NVJ) through Nvj1 (Pan et al., 2000; H et al., 2017) and the recently described vacuole-lipid droplet contact site (vCLIP) through Ldo45/16 (Álvarez-Guerra et al., 2024; Diep et al., 2024). In addition, Vac8 contributes to autophagosome biogenesis at the vacuolar membrane through recruitment of the PIK3C3 complex (Hollenstein et al., 2019; Gatica et al., 2021; Hollenstein et al., 2021; Lei et al., 2021).

**Figure 5.**
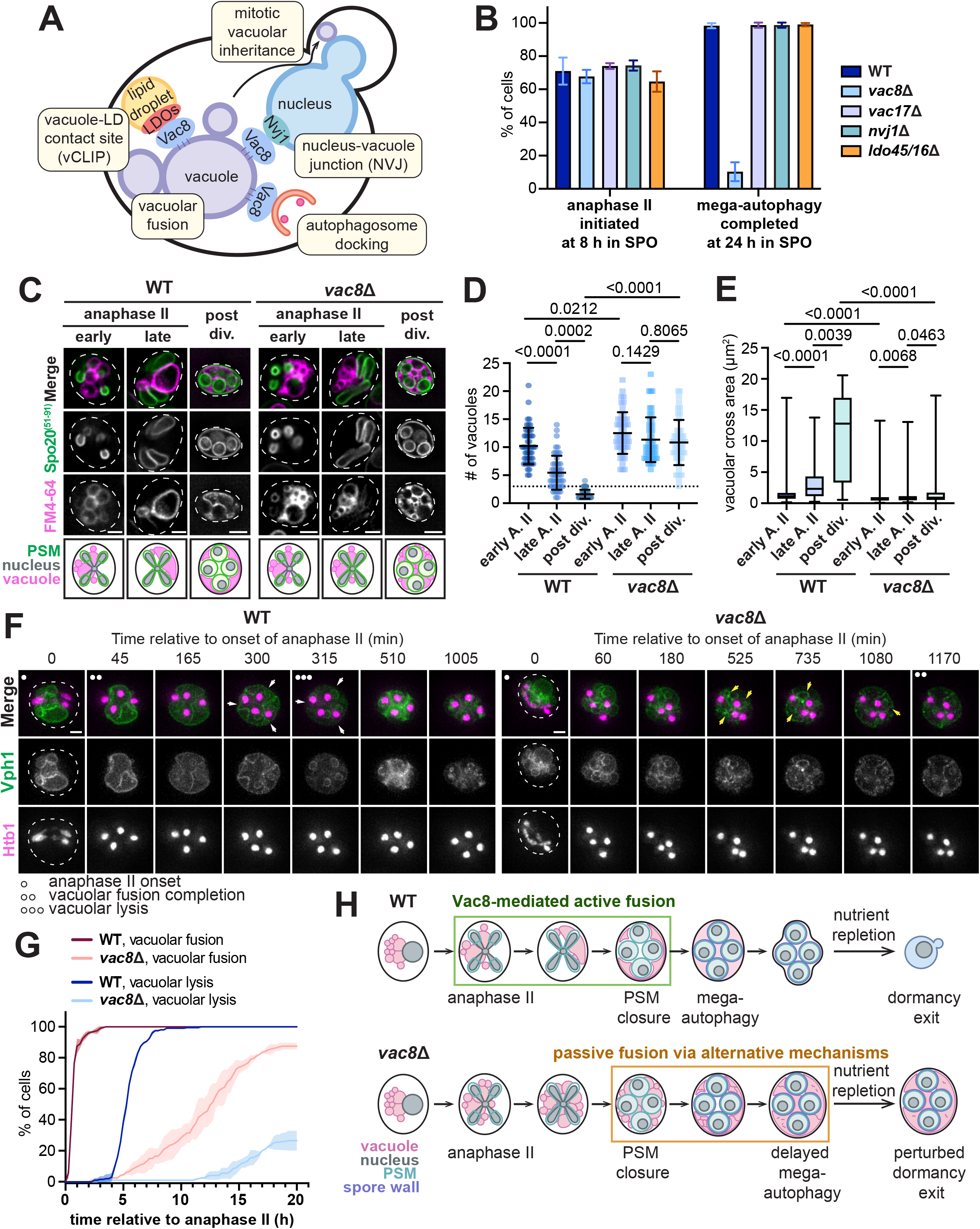
Vac8-mediated vacuolar fusion during anaphase II primes post-meiotic vacuolar lysis. **(A)** Schematic of Vac8-associated pathways identified in mitotic cells. Vac8 mediates mitotic vacuolar inheritance and forms membrane contacts with both the nucleus and the lipid droplets. Vac8 also confines autophagosome assembly on vacuoles and is required for vacuolar fusion. **(B)** The percentage of WT and indicated mutants that had initiated anaphase II at 8 h in SPO and that had lysed their precursor-cell vacuoles at 24 h in SPO. Error bars = SD of three replicates, n = 100 cells per replicate. **(C)** Widefield microscopy images of WT and *vac8*Δ cells progressing through anaphase II. Precursor vacuoles were visualized by FM4-64 staining prior to switching to SPO. The prospore membrane (PSM) was marked by GFP-SPO20^51-91^. The stages at anaphase II were determined by PSM morphology. Post-division (post div.) samples were harvested at 9.5 h in SPO. Image represents a single z-slice. Scale bar = 2 µm. **(D and E)** Quantification of (C) showing the number of vacuoles per cell (D) and the cross-section areas of individual vacuoles (E) in WT and *vac8*Δ cells at indicated stages during gametogenesis. A. II, anaphase II. The dashed line in (D) indicates vacuole number = 3 per cell. For statistical analysis information, see Table S5. The p-values are indicated in the graph. **(F)** Montage of WT and *vac8*Δ cells expressing Vph1-GFP progressing through gametogenesis. Chromatin was visualized with Htb1-mCherry. The time point depicting anaphase II onset was defined as 0 min and indicated by the single white dot on the top left corner of the merged image. The time point depicting vacuolar fusion completion (defined by less than 3 vacuoles per cell) was indicated by the double white dots on the top left corner of the merged image. White arrows indicate the disappearance of Vph1 membrane signal, and the time point depicting the lysis of precursor vacuoles was indicated by the triple white dots on the top left corner of the merged image. Yellow arrows indicate the membrane remnants formed during swelling-mediated vacuolar fusion. Images are shown as MIPs of z-stacks. Scale bar = 2 µm. **(G)** The percentage of WT and *vac8*Δ cells completed vacuolar fusion or vacuolar lysis plotted relative to the time of anaphase II onset for each cell. Vacuolar fusion completion was determined by whether the cell contained less than 3 precursor vacuoles, and vacuolar lysis was marked by the disappearance of Vph1 membrane signal on the precursor vacuoles. Shaded region indicates SD of three replicates, n = 100 cells per replicate. **(H)** Schematic of the meiotic vacuolar remodeling pathway. During meiosis II, precursor-cell vacuoles start to fuse in a Vac8-dependent manner. After PSMs close, precursor vacuoles continue fusing and swelling, ultimately forming a single vacuole that fills most of the ascal space unoccupied by spores. After spores fully developed spore walls, the mega-autophagy is triggered, rupturing the swollen precursor vacuole and degrading all uninherited contents remaining in the ascus. Upon nutrient repletion, spores in the asci that have completed mega-autophagy are able to exit from dormancy and re-enter mitosis. In the absence of Vac8, vacuoles fail to actively fuse during late stages of meiosis II. Instead, the precursor-cell vacuoles gradually swell and force only intermittently much later, leading to a delay in mega-autophagy. Without execution of mega-autophagy, spores in the asci were unable to efficiently germinate in response to nutrient replenishment. Vacuoles are indicated in light purple, the nucleus is indicated in gray, the PSM is indicated in light blue, and the spore wall is indicated in indigo.

We first tested whether Vac8-dependent tethering to myosin, nuclear envelope or lipid droplets contributes to mega-autophagy by disrupting the corresponding Vac8-interacting proteins. Using GFP-tagged Vph1to visualize vacuolar membranes, we found that deletion of *VAC17*, *NVJ1*, or *LDO45/16* impaired neither meiotic progression nor post-meiotic vacuolar lysis (Fig. 5 B; Fig. S3A). Thus, Vac8 promotes mega-autophagy independently of its established functions in vacuolar inheritance and vacuolar contacts with the nuclear envelope or lipid droplets.

We next asked whether the mega-autophagy defect could instead arise from the established role of Vac8 in autophagosome biogenesis (Hollenstein et al., 2019; Gatica et al., 2021; Hollenstein et al., 2021; Lei et al., 2021). Because direct disruption of autophagosome formation severely impairs sporulation, we assessed whether *VAC8* deletion itself compromises autophagosome-mediated degradation during meiosis. It has been shown that autophagosome-based degradation of organelles occurs during meiosis (Otto et al., 2021). Using GFP-tagged Atg8 as an autophagosome marker, we observed comparable free-GFP accumulation in WT and *vac8*Δ cells during early meiosis, indicating that bulk autophagy remained largely intact in the absence of Vac8 (Fig. S3B). At 12 h in SPO, WT cells exhibited a pronounced additional increase in free GFP associated with mega-autophagy, which was absent in *vac8Δ* cells (Fig. S3B). Thus, loss of Vac8 selectively impairs post-meiotic mega-autophagy without broadly disrupting autophagosome-mediated degradation during meiosis. Together, these findings suggest that Vac8 performs a distinct function during meiotic development that prepares precursor-cell vacuoles for their subsequent post-meiotic destruction.

### Vac8-facilitated vacuolar fusion during anaphase II primes mega-autophagy

We next sought to identify the meiotic function of Vac8 that enables mega-autophagy. Vac8 promotes homotypic vacuolar fusion in mitotically growing cells (Wang et al., 2001). Notably, post-meiotic cells contain characteristically large, fused vacuoles prior to their rupture (Roeder and Shaw, 1996; Eastwood and Meneghini, 2015). We thus asked whether Vac8 promotes developmentally regulated vacuolar fusion during meiosis.

Using FM4-64 to follow precursor-cell vacuoles together with GFP-Spo20^51-91^ to monitor PSM development, we observed a striking remodeling of the vacuolar compartment during anaphase II (Fig. 5 C-E). At early anaphase II, when PSMs had only begun to form, WT cells contained multiple small precursor vacuoles. As PSMs elongated around the gamete nuclei, these vacuoles progressively fused into fewer, larger structures, ultimately forming a single enlarged vacuole following PSM enclosure (Fig. 5 C-E). In comparison, the vacuolar fusion during late meiosis appears to be absent in *vac8*Δ cells (Fig. 5 C-E), with no significant change in the number of vacuoles per cell over time. While vacuole size increases slightly in *vac8*Δ cells during anaphase II, the post-division size is dramatically smaller than in WT cells (Fig. 5E). These results indicate that Vac8 is required for meiotic vacuolar fusion. In support of a direct role in this process, Vac8-GFP became enriched at the interface between contacting vacuoles during meiosis II (Fig. S3 C and D). Vac8 enrichment at vacuole–vacuole contact sites was significantly greater than that observed for the general vacuolar membrane marker Vph1 (Fig. S3, C and D), indicating that the increased Vac8 signal does not simply reflect local membrane accumulation and instead suggests preferential recruitment of Vac8 to sites of vacuolar fusion.

To resolve the dynamics of this remodeling process, we followed Vph1-GFP in WT and *vac8*Δ cells by time-lapse imaging (Fig. 5F). Nearly all WT cells consolidated their vacuolar compartment to fewer than three vacuoles within 2 h of anaphase II onset (Fig. 5, F and G). In contrast, vacuolar fusion was markedly delayed in *vac8*Δ cells (Fig. 5, F and G). Precursor vacuoles nevertheless began to swell after anaphase II in both WT and *vac8*Δ cells, indicating that vacuolar swelling occurs independently of Vac8 (Fig. 5 E and F). In *vac8*Δ cells, this progressive swelling was eventually accompanied by slower vacuolar fusion, suggesting the existence of a secondary, Vac8-independent route to vacuolar coalescence (Fig. 5, F and G).

The timing of Vac8-dependent vacuolar fusion preceded mega-autophagy. In WT cells, precursor vacuoles underwent lysis approximately 4-5 h after completion of vacuolar fusion, whereas only ∼30% of *vac8Δ* cells completed vacuolar lysis by the end of the imaging period (28 h in SPO; Fig. 5, F and G). Together with our finding that Vac8 is specifically required before anaphase II to support subsequent mega-autophagy (Fig. 3, C–G), these results identify rapid Vac8-dependent vacuolar fusion during anaphase II as a developmental remodeling event that primes precursor-cell vacuoles for their later post-meiotic lysis (Fig. 3H).

Finally, we asked whether other components known to promote homotypic vacuolar fusion during vegetative growth are similarly required for this meiotic remodeling event. We examined components of the HOPS (homotypic fusion and protein sorting) complex, the SNARE machinery (Wickner, 2002; Krämer and Ungermann, 2011), the vacuolar V-ATPase membrane component Vph1 (Bayer et al., 2003), and the vacuolar transporter chaperone protein Vtc3 (Müller et al., 2002). Several HOPS and SNARE mutants (*ypt7*Δ, *sso1*Δ, *vam3*Δ, and *vam7*Δ) failed to sporulate and therefore could not be evaluated for mega-autophagy. For remaining mutants that were able to sporulate, no detectable defects were identified in meiotic vacuolar fusion or post-meiotic vacuolar lysis (Fig. S3, E–H). Although mutants lacking the vacuolar SNARE component Nvy1 showed more severe vacuolar fragmentation at early anaphase II than *vac8*Δ cells, >80% of *nyv1*Δ cells completed vacuolar fusion at 9.5 h in SPO and did not show mega-autophagy defects at 24 h in SPO (Fig. S3, E, G, and H). These findings further distinguish the meiotic remodeling program from canonical vacuolar fusion pathways and point to a specialized role for Vac8 in reorganizing precursor-cell vacuoles following chromosome segregation in anaphase II.

In this study, we identify Vac8 as a key regulator of meiotic vacuolar remodeling. Following chromosome segregation in anaphase II, Vac8 promotes the rapid fusion of precursor-cell vacuoles, which primes the resulting vacuole for post-meiotic mega-autophagy. In the absence of Vac8, this developmentally-timed fusion event is impaired, although vacuoles eventually coalesce more slowly, potentially as a consequence of post-meiotic vacuolar swelling. This delayed remodeling is associated with severe defects in mega-autophagy and, ultimately, the ability of intact spores to exit dormancy upon nutrient repletion (Fig. 5H). Together, our findings reveal a developmentally regulated sequence in which Vac8-dependent vacuolar fusion during meiosis II prepares the precursor vacuole for its subsequent destruction after meiosis, thereby promoting efficient spore germination.

How does Vac8 facilitate vacuolar fusion? Even in mitotically dividing cells, the mechanism by which it promotes vacuolar fusion remains unclear. Structural studies have shown that Vac8 can form homodimers (Kim et al., 2023), raising the possibility that Vac8 self-association contributes to vacuolar fusion. Consistent with this possibility, we observed enrichment of Vac8 at vacuole-vacuole contact sites during anaphase II (Fig. S3, C and D). However, Vac8 dimerization is also required for interactions with its several binding partners (Kim et al., 2023), making it difficult to isolate a specific role for dimerization in vacuolar fusion. Similarly, previous structure-function analyses using deletions of individual Vac8 ARM domains found that domains required for vacuolar fusion also contribute to other Vac8 functions, thereby precluding the generation of clean separation-of-function mutants. (Wang et al., 1998). Notably, *VAC8* deletion does not disrupt the formation of vacuole-vacuole contacts in both mitotic (Wang et al., 2001; Fig. S3F) and meiotic conditions (Fig. 5C; Fig. S3, C-F). These observations place Vac8 downstream of vacuole-vacuole contact formation and suggest that it promotes a subsequent step leading to membrane fusion. In situ structural characterization of vacuole-vacuole interfaces in WT and *vac8*Δ cells may therefore reveal how Vac8 facilitates the fusion of adjacent vacuolar membranes.

Our findings suggest that Vac8-mediated vacuolar fusion during anaphase II primes the precursor vacuole for mega-autophagy, raising the question of what subsequently triggers vacuolar membrane rupture. One possibility is that meiotic changes in vacuolar lipid composition render the vacuolar membrane susceptible to rupture. In addition, precursor vacuoles undergo pronounced swelling prior to rupture independently of Vac8 (Fig. 5 E and F), suggesting that osmotic forces may contribute to destabilizing the vacuolar membrane. The trigger for mega-autophagy may also be coupled to gamete maturation, because mega-autophagy requires spore enclosure and spore wall development (Eastwood and Meneghini, 2015). Such a signal that triggers vacuolar rupture could arise from molecules associated with the developing spore surface or from changes in the physical or chemical environment accompanying spore maturation. Notably, *vac8*Δ mutants show no apparent defects in spore wall formation (Fig. S2 A and B), supporting the notion that Vac8-dependent meiotic vacuolar remodeling is not coupled to spore wall formation in triggering mega-autophagy.

Completion of mega-autophagy is required for spores within their native environment to efficiently exit dormancy and re-enter mitotic growth upon nutrient repletion (Fig. 4). Intriguingly, this defect does not appear to reflect an intrinsic inability of spores to resume growth, as enzymatic removal of the ascus wall with zymolyase largely restores their response to nutrient-rich conditions (Fig. 4). These findings suggest that, in the absence of mega-autophagy, the persistence of material within the precursor-cell cytoplasm creates an environment that is unfavorable for enclosed spores to germinate. Additionally, degradation of uninherited cellular material could generate building blocks and/or metabolites that are recycled by developing or dormant spores, potentially supporting survival during prolonged nutrient limitation. Determining whether and how spores access such molecules will further define the physiological benefits of mega-autophagy. Interestingly, the mammalian Vac8 ortholog ARMC3 is predominantly expressed in reproductive tissues and is required for male fertility (Pausch et al., 2016; Lei et al., 2021; Rahim et al., 2024). During mouse spermatogenesis, ARMC3 promotes ribophagy, which has been proposed to support mitochondrial metabolism in developing sperm (Lei et al., 2021). Together with our finding that Vac8-dependent mega-autophagy promotes efficient spore germination, these observations raise the possibility that Vac8-family proteins have evolved related roles in coupling cellular remodeling to gamete fitness across eukaryotes.

## Materials and methods

### Reagents

Plasmids, chemicals, and antibodies used in this study are listed in Table S1.

### Yeast strains, plasmids, and primers

A complete list of all yeast strains, plasmids, and primers in this study can be found in Table S2, Table S1, and Table S3, respectively.

All *S. cerevisiae* strains in this study are derivatives of SK1 (Kane and Roth, 1974) and are homozygous diploids unless specified. The following alleles were constructed in previous studies: *HTB1-mCherry* (Matos et al., 2008); *pATG8-nucGEM* (Ruediger et al., 2025); *pATG8-GFP-SPO20^51-91^ VPH1-GFP*, *NUP170-GFP*, and *pCUP1-OsTIR^F74G^* (King et al., 2019); *CIT1-*GFP (Sawyer et al., 2019); *SEC63-GFP* and *GFP-ATG8* (Otto et al., 2021); *RPL26B-GFP* and *RPL29-GFP* (Eisenberg et al., 2018). The following used alleles were introduced in this study: *vac8*Δ, *PMA1-GFP*, *VAC8-3V5-IAA17*, *vac17*Δ, *nvj1*Δ, *ldo45/16*Δ, *dit1*Δ, *dit2*Δ, *VAC8-GFP*, *vps33*Δ, *nyv1*Δ, *vph1*Δ, and *vtc3*Δ. Specifically, heterozygous strain containing the *PMA1-GFP* allele was used to minimize potential functional deficiency caused by tagging. The *VAC8-3V5-IAA17* was created by inserting a 3V5-IAA17 fragment (King et al., 2023) at the C-terminus of Vac8 in *flo8*Δ strains. Depletion experiments were also performed in *FLO8+* strains and showed similar results. Only data from *flo8*Δ strains are shown here, as these cells pelleted more efficiently during washing steps, minimizing cell loss. For the non-depleted control, a *URA3* cassette (Youk and Lim, 2014) was integrated at the *URA3* locus to match the auxotrophic background of the OsTIR1 group, as the *pCup-OsTIR(F74G)* carries a *URA3* marker. For *ldo45/16*Δ allele, the entire *LDO16* ORF is localized within the *LDO45* ORF, and thus deleting the *LDO45* ORF leads to the deletion of both *LDO45* and *LDO16* (Álvarez-Guerra et al., 2024; Diep et al., 2024).

Deletions and endogenous C-terminal tagging of genes were achieved by standard PCR-mediated methods with the indicated selection cassette listed in Table S2 and oligo pairs listed in Table S3. Plasmids used for amplification are listed in Table S1 (Janke et al., 2004; Sheff and Thorn, 2004; Ruediger et al., 2025; Powers et al., 2022). Strains expressing *pATG8-nucGEM*, *pATG8-GFP-SPO20^51-91^*, *VPH1-GFP*, or *GFP-ATG8* were created by integrating a single-integration vector at the *TRP1*, *LEU2*, *HIS3*, or *URA3* locus, respectively. For single-copy integration, plasmids were digested by PmeI for linearization, followed by standard transformation techniques.

Yeast transformation was performed in strain 13 (SK1 wild-type haploid, *MATa*) using the lithium acetate method (ref). Proper deletion, integration, or tagging was verified by PCR, followed by fluorophore visualization, Western blot, or sequencing, where applicable. Each haploid strain carrying gene deletion, integration, and/or endogenous tagging was then backcrossed to strain 14 (SK1 wild-type haploid, *MATα*), and the resulting heterozygous diploids were sporulated and dissected to obtain genetically modified strains of both mating types. If applicable, haploid strain containing required markers and alleles were crossed in, followed by dissection to obtain both mating types. Homozygous diploids were then generated by crossing haploid strains of opposite mating types carrying the same genotypes.

### Yeast cell culture and media

For experiments conducted during vegetative growth, cells were grown in YPD (1% yeast extract, 2% peptone, 2% glucose, 22.4 mg/L uracil, and 80 mg/L tryptophan) for 20–24 h at room temperature to reach a cell density of OD_600_ ≥10. Cultures were then back-diluted to OD_600_ = 0.1 and allowed to grow for 4–5 h at 30°C. For imaging experiments, cells were examined at a density of OD_600_ = 0.6–0.8.

For sporulation, diploid cells were first grown in YPD for around 20–24 h at room temperature to reach an OD_600_ ≥10. The cultures were then diluted in BYTA (1% yeast extract, 2% Bacto tryptone, 1% potassium acetate, and 50 mM potassium phthalate) to OD_600_ = 0.25 and grown for 12–16 h at 30°C. After reaching an OD_600_ = 5–8, the cells were pelleted (1,900 × *g*, 2 min), washed once in sterile MilliQ water, and resuspended in the sporulation media SPO (2% potassium acetate, 40 mg/L adenine, 40 mg/L uracil, 10 mg/L histidine, 10 mg/L leucine, and 10 mg/L tryptophan, pH adjusted to 7.0) to OD_600_ = 1.85. Meiotic cultures were shaken at 30°C for the duration of the experiment. At all stages, the flask size was 10 times the culture volume to ensure proper aeration.

For experiments using the *VAC8-3V5-IAA17* allele, CuSO_4_ was added to a final concentration of 50 µM at the indicated times to induce expression of pCUP1-OsTIR1^F74G^ followed immediately by 100 µM 5-Ph-IAA (Sigma-Aldrich, SML3574).

### Sporulation efficiency, spore viability, and germination

Sporulation efficiency was assessed using cell culture sporulated for 24 h in SPO. Cells contained Htb1-mCherry, and portions of progenitors forming 0 spores (unsporulated), 1-2 spores (monads/dyads), or 3-4 spores (triads/tetrads) were scored using fluorescence microscopy. Spore viability was assessed by the growth of sporulated asci or individual spores on YPD agar plates. For whole asci, 50 µl of yeast cultures that were sporulated for 24 h were diluted 1:10 in SPO and placed onto YPD agar plates. Each ascus was placed at specific coordinate under a dissection scope. For individual spores, 50 µl of yeast cultures that were sporulated for 24 h were harvested, pelleted, and treated with 1 mg/mL zymolyase in 15 µl MiliQ water (MP Biomedicals, 083209-CF) at room temperature for 7 minutes. The cells were then resuspended in 500 µl SPO and sonicated for 5 pulses to better separate spores. Individual spores were placed at specific coordinate under a dissection scope. For each pair of WT and *vac8*Δ cells, the dissection was finished within the same hour to minimize time difference. After growing at 30°C for 48 h, plates were imaged by Gel Doc XR+ Gel Documentation System (BioRad) with colonies facing up, and colony size was quantified in FIJI (RRID:SCR_002285, Schindelin et al., 2012) by image thresholding and particle size analysis. Germination of ascus-encapsulated or individual spores was assessed by time-lapse microscopy, and details are described in the live-cell imaging session below.

### Vacuolar staining by FM4-64

For vegetatively growing cells, FM4-64 staining was modified from Amberg et al., 2006. Cells were grown in YPD for 20–24 h at room temperature to reach a cell density of OD_600_ ≥10. Cultures were then back-diluted to OD_600_ = 0.1 and allowed to grow for 4–5 h at 30°C to reach a density of OD_600_ = 0.6–0.8. For staining, FM™ 4-64 Dye (N-(3-Triethylammoniumpropyl)-4-(6-(4- (Diethylamino) Phenyl) Hexatrienyl) Pyridinium Dibromide) (FM4-64; Invitrogen™, T13320) was added to 2 ml of cells to a final concentration of 1 μg/ml. After incubation at 30°C in the dark for 1 h, cells were washed twice with 1 ml YPD and resuspended in 1 ml YPD to allow incubation for another hour. Cells were harvested and resuspended in 50 μl of imaging buffer (10 mM HEPES, pH 7.6, and 5% glucose) prior to imaging.

Staining of sporulating cells with FM4-64 was modified from Roeder and Shaw, 1996. Diploid cells were first grown in YPD for around 20–24 h at room temperature to reach an OD_600_ ≥10. The cultures were then diluted in BYTA to OD_600_ = 0.25 and grown for 12–16 h at 30°C. Before transferring to SPO media, FM4-64 was added to BYTA culture to a final concentration of 1 μg/ml. After incubation at 30°C in the dark for 1 h, cells were pelleted (1,900 × *g*, 2 min), washed once in sterile MilliQ water, and resuspended in the sporulation media SPO. Cells were incubated at 30°C in the dark for sporulation until imaging. For imaging, 500 μl of cell cultures that had been sporulated for indicated time were harvested by centrifugation (1,900 × *g*, 2 min), and then 450 μl of supernatant was discarded. The rest of 50 μl culture were used to resuspend the cells. Images were obtained at room temperature using the TRITC filter set. Detailed imaging conditions are indicated in Table S4.

### Dityrosine autofluorescence examination

Dityrosine fluorescence assay were performed as previously described (Suda et al., 2009b). Briefly, 50 μl of cells that had been sporulated for 24 h in SPO were pelleted and resuspended in 50 μl of either conditioned SPO or 5% NH_4_OH (Thermo Scientific Chemicals, AC458690010).

Imaging conditions are indicated in Table S4. The dityrosine autofluorescence intensity was quantified by drawing a line scan across the periphery of each sporulated ascus and subtracting the background intensity from the peak intensity.

### Fluorescence microscopy

Images were acquired using a DeltaVision Elite wide-field fluorescence microscope (GE Healthcare) and a PCO Edge scientific complementary metal–oxide–semiconductor camera, with softWoRx software and a 60× NA1.42 oil-immersion Plan Apochromat objective. GFP and RFP filter sets were used, with the setting information for acquiring each figure panel provided in Table S4. Images were deconvolved using softWoRx software (GE Healthcare) and 3D iterative constrained deconvolution with 15 iterations and enhanced ratio. Whether images were single-z plane or maximum intensity z-projected over the range of acquisition in FIJI (RRID:SCR_002285, Schindelin et al., 2012) are indicated in Table S4.

### Live-cell imaging

For still images, 1 × 10^7^ of cells were obtained from vegetatively growing or SPO culture. Cells were sonicated for 5 pulses (Branson; 100-132-1623R & 101-135-066R) and harvested by centrifugation (1,900 × *g*, 2 min). For vegetatively growing culture, cells were washed once in MiliQ water and resuspended in 50 μl imaging buffer (10 mM HEPES, pH 7.6, and 5% glucose) prior to imaging. For SPO cultures, 450 μl of supernatant was discarded, and the remaining 50 μl supernatant was used to resuspend the cells. Images were obtained at room temperature.

For time-lapse imaging, cells were imaged in an environmental chamber heated to 30°C, using either the CellASIC ONIX Microfluidic Platform or glass-bottom 96-well plates. For the CellASIC ONIX Microfluidic Platform (EMDMillipore, Y04D microfluidics plates), cells that had been sporulated for 3–5 h in SPO were sonicated for 5 pulses and transferred to the microfluidic plates and loaded at 8 psi for 5 s. Conditioned SPO was prepared by filtering out WT cells from SPO culture that had been sporulated for 5 h and was applied with a constant flow rate pressure of 2 psi for 15–20 h. For 96-well plates, each well to be imaged was precoated with 1 mg/ml concanavalin A for 30 min, washed once with MiliQ water, and then dried completely before adding cells. For imaging during sporulation, 20 μl of cells that had been sporulated for 3–5 h in SPO were sonicated for 5 pulses and added to each well. After cells were adhered for 15 min, the remaining culture was then removed to get rid of extra cells, and 100 μl of conditioned SPO was added to each well. Images were acquired every 15 min for 18–20 h. For whole-asci germination experiments, cell culture that had been sporulated for 24 h were sonicated for 5 pulses and diluted 1:5 before adding to each well. For the germination experiments of individual spores, 500 μl of cell culture that had been sporulated for 24 h were harvested and pelleted. The pellets were then treated with 1 mg/ml zymolyase (MP Biomedicals) for 7 min and resuspended in 500 μl MiliQ water. The resuspended cells were then sonicated for 6 pulses twice and diluted 1:10 before adding to each well. After asci/spores were adhered for 15 min, the remaining cell cultures were then removed to get rid of extra cells, and 100 μl of YPD was added to each well. for sporulating culture. Images were acquired every 5 min for 10 h.

### Image quantification and statistical analysis

Anaphase II entry was determined by the timing of the division of four gamete nuclei. For the clearance of excluded substrates, the timing of excluded nucGEM, Nup170-GFP, Cit1-GFP, Sec63-GFP, or Pma1-GFP clearance was scored in cells that were sporulated and formed spores that were visible by brightfield. Substrate clearance was defined as the whether the excluded fluorescence signal was no longer visible. For nucGEM sequestration experiments, sequestration was defined as the aggregate no longer associating with chromatin on any focal plane after the four anaphase II nuclei became distinct. Nucleoporin sequestration was defined as the enrichment around the compartment that was not associated with any gamete nucleus after the four anaphase II nuclei became distinct.

For vacuolar lysis experiments, the timing of vacuolar membrane disruption was scored in cells that sporulated and formed spores that were visible under brightfield. Mega-autophagy completion was defined as whether Vph1-GFP or FM4-64 membrane signal became invisible and rather diffused in the ascus. Occasionally, the asci appeared to collapse prior to vacuolar membrane disappearance. Since we were unable to interpret these changes in vacuolar morphology, these cells were not included in our quantification. For vacuolar fusion during meiosis, stages of anaphase II were determined based on PSM size indicated by GFP-SPO20^51-91^. Vacuolar fusion was determined by whether the sporulating cell only contains less than 3 precursor-cell vacuoles.

For germination of in-ascus or individual spores, the timing of dormancy exit was defined by the appearance of the first bud after YPD addition. For spores in asci, only the first bud generated from the entire ascus was recorded. Ribosomal protein intensity was quantified as previously described (Eisenberg et al., 2018). Mean intensity of signal from the cells or spores was measured in FIJI (RRID:SCR_002285, Schindelin et al., 2012) and was divided by the background signal from the same image for normalization. For GFP enrichment at vacuole-vacuole contact sites, GFP intensity was quantified by detecting the line profiles along the contacting and non-contacting regions of the vacuolar membrane separately, and the mean intensity along each line was measured in FIJI (RRID:SCR_002285, Schindelin et al., 2012). Contact-site intensity was then normalized to that of the non-contacting membrane. The 3D plot was created in FIJI, with the x- and y-axes indicating the image dimensions in microns, and the z-axis indicating fluorescence intensity (a.u.).

All statistical analysis was performed in Prism (GraphPad), and the statistical test used is indicated in the corresponding figure legend and listed in Table S5.

### Immunoblotting

Immunoblotting was performed similarly as previously described (King et al., 2019). Briefly, 2.2 OD units of cells were harvested, resuspended in 1 ml of 5% trichloroacetic acid, incubated overnight at 4°C. The cells were subsequently washed with 750 μl 10 mM Tris pH 8.0 and then 750 μl of acetone, before being left to dry overnight. Cell pellets were then resuspended in 100 μl of lysis buffer (50 mM Tris-HCl pH 8.0, 1 mM EDTA, 15 mM Tris pH 9.5, 3 mM DTT, 1X cOmplete EDTA-free inhibitor cocktail [Roche]) and pulverized with glassbeads using a Mini-Beadbeater-96 (BioSpec). 50 μl of 3X SDS sample buffer (187.5 mM Tris pH 6.8, 6% β-mercaptoethanol, 30% glycerol, 9% SDS, 0.05% bromophenol blue) were then added to each sample and boiled at 95°C for 5 min.

Extracted proteins were separated by polyacrylamide gel electrophoresis using 4–12% Bis-Tris Bolt gels (Thermo Fisher) and transferred onto nitrocellulose membranes (0.45 μm, Bio-rad) using a semi-dry transfer apparatus (Trans-Blot Turbo Transfer System, Bio-Rad). The membranes were blocked with Odyssey PBS Blocking Buffer (LICORbio) at room temperature for 1 h. The blots were incubated overnight at 4°C with a mouse anti-GFP antibody (RRID: AB_2313808, 632381, Takara) or a mouse anti-V5 antibody (RRID: AB_2556564; 461157, Invitrogen) at a 1:2000 dilution in blocking buffer. As a loading control, we monitored Hxk2 levels using a rabbit anti-hexokinase antibody (RRID: AB_219918, 100–4159, Invitrogen) at a 1:20,000 dilution in blocking buffer. Membranes were washed in PBST (1 × PBS with 0.1% Tween-20) and incubated with an anti-mouse secondary antibody conjugated to IRDye 800CW at a 1:15,000 dilution (RRID: AB_621842, 926–32210, LICORbio) and an anti-rabbit antibody conjugated to IRDye 680RD at a 1:15,000 dilution (RRID: AB_10956166, 926–68021, LICORbio) to detect the GFP or V5 epitope and Hxk2, respectively. Immunoblot images were generated using the Odyssey CLx system (LICORbio). For quantification, band intensity was analyzed using ImageStudio software (LICORbio).

### Online supplemental material

Fig. S1 shows that deleting *VAC8* does not perturb meiosis entry or GUNC sequestration in SK1 cells, but delays the post-meiotic degradation of excluded uninherited organellar contents. Fig. S2 contains data suggesting that *VAC8* deletion does not inhibit spore wall development, that *vac8Δ* spores exhibit delayed germination and ribosomal protein expression, and that Vac8 is expressed throughout gametogenesis and can be depleted via the auxin-degron system. Fig. S3 shows that several known Vac8 interactors and vacuolar fusion pathway components are dispensable for mega-autophagy, and that Vac8 is enriched at the vacuole-vacuole junction during anaphase II. Table S1 lists the chemicals, antibodies, plasmids, and software used in this study. Table S2 lists the yeast strains used in this study. Table S3 lists the oligonucleotides used in this study. Table S4 includes the imaging conditions of all microscopy images shown in this study. Table S5 lists the quantification and statistical analysis information related to this study.

## Data availability

This paper does not report original code. All reagents used in this study are available upon request. The data underlying all figures are available in the published article and supplemental information, and from the corresponding author.

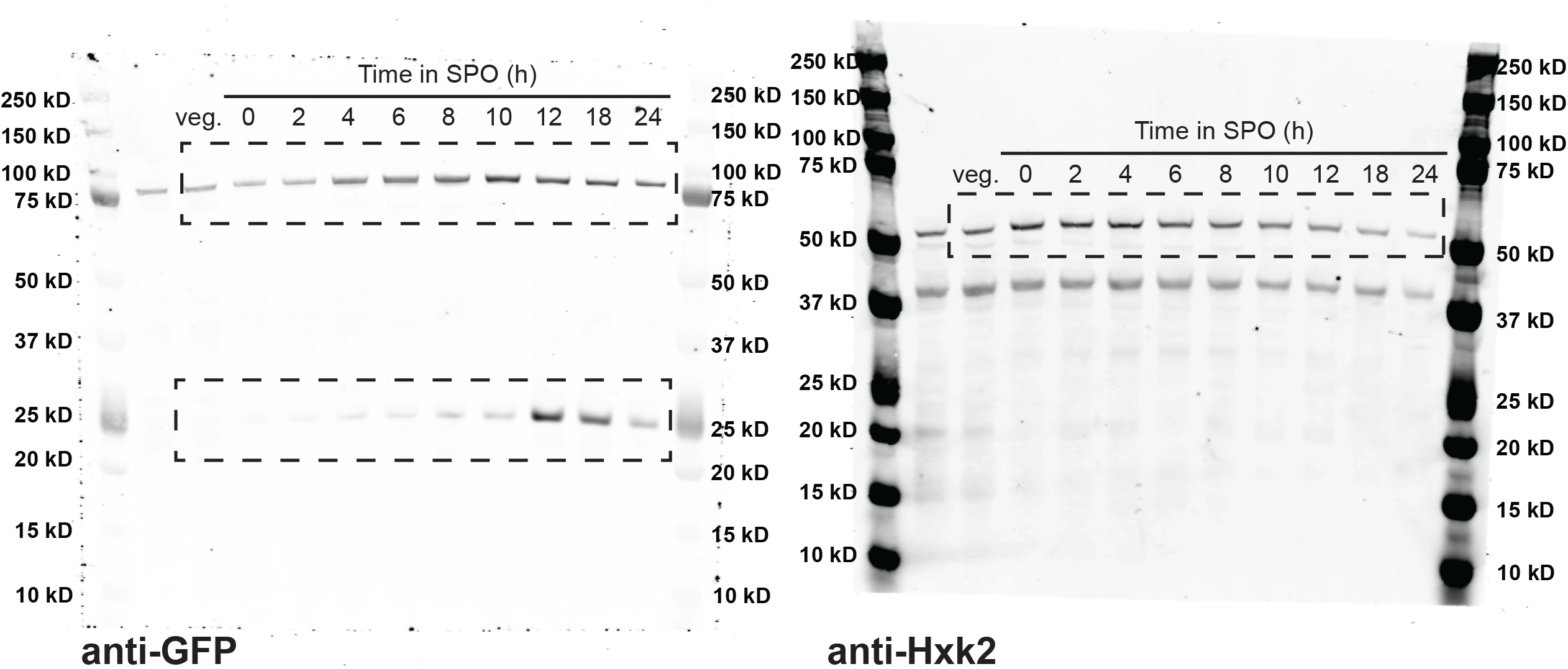
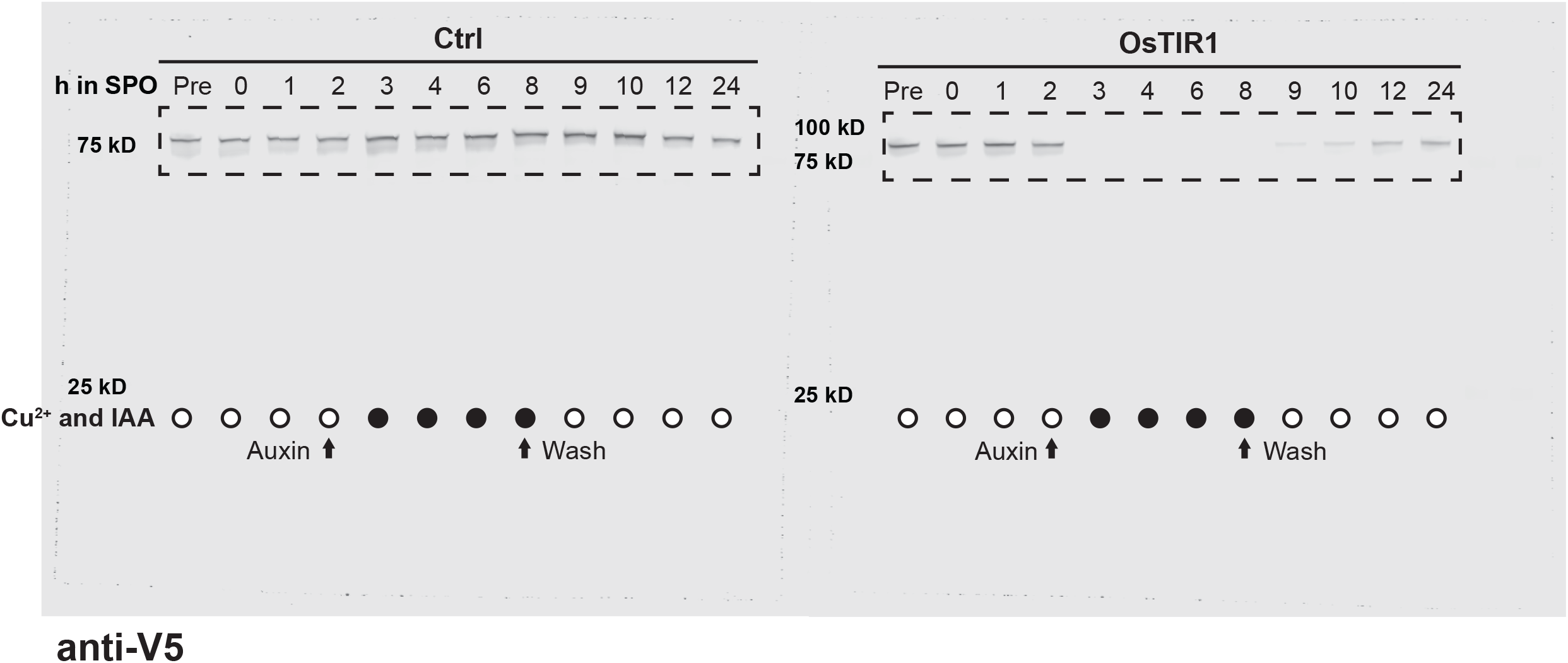
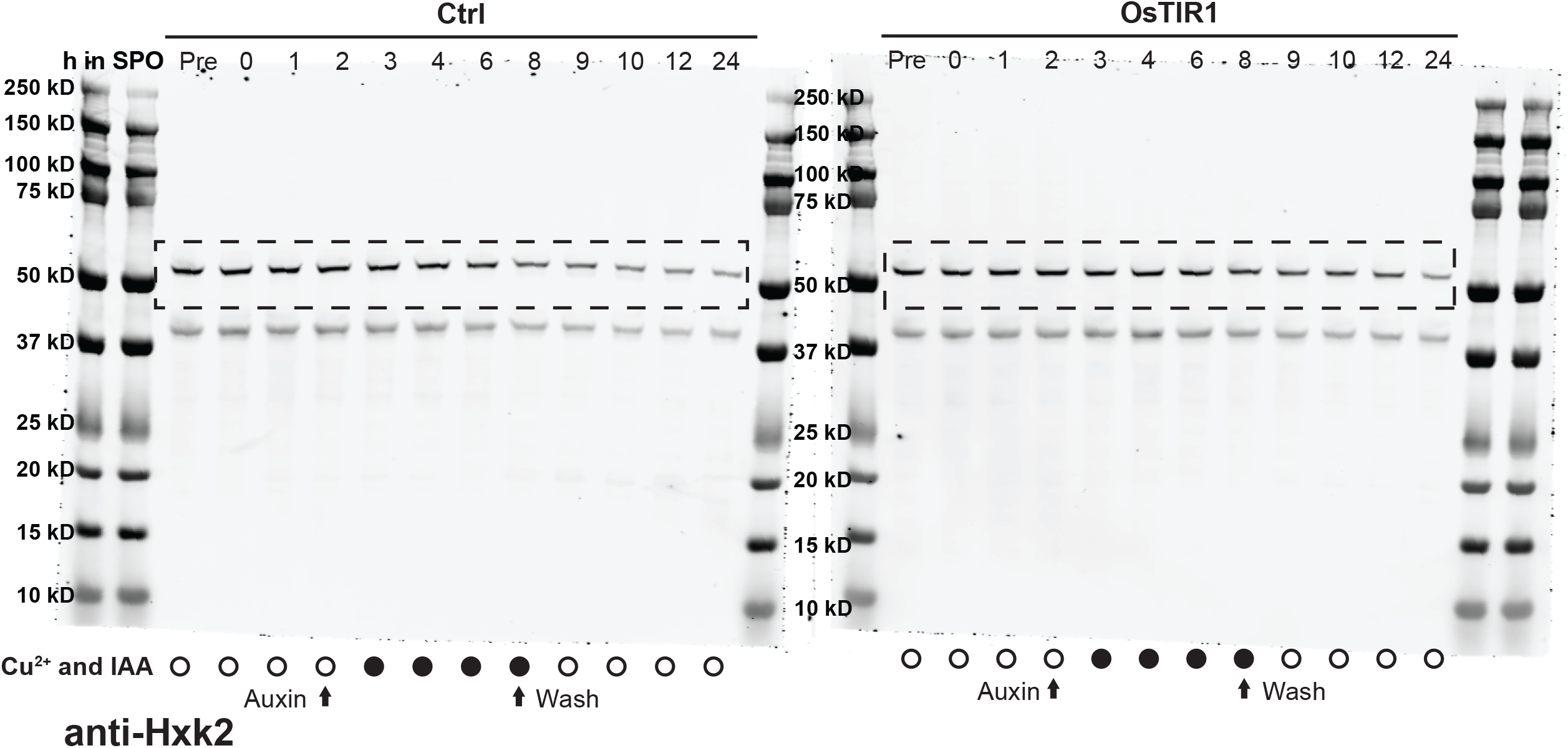

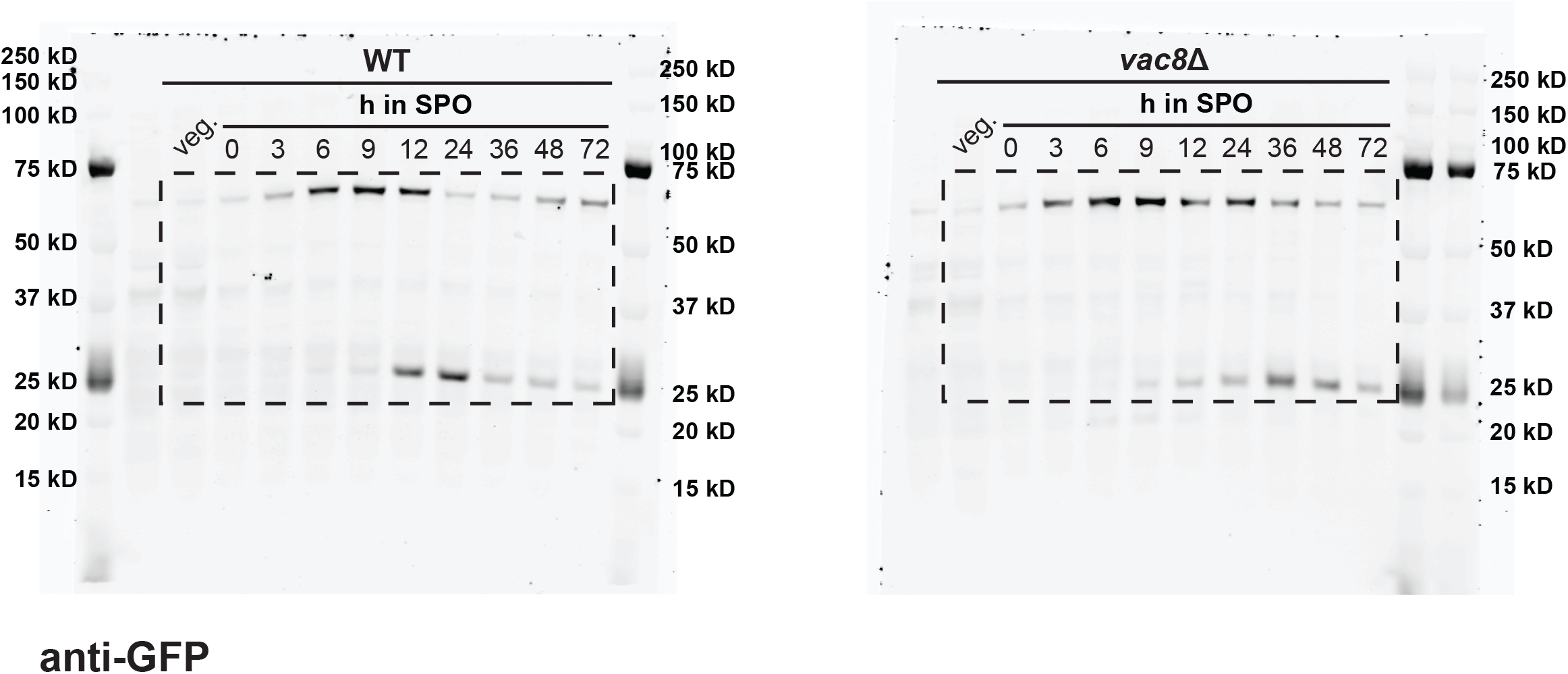
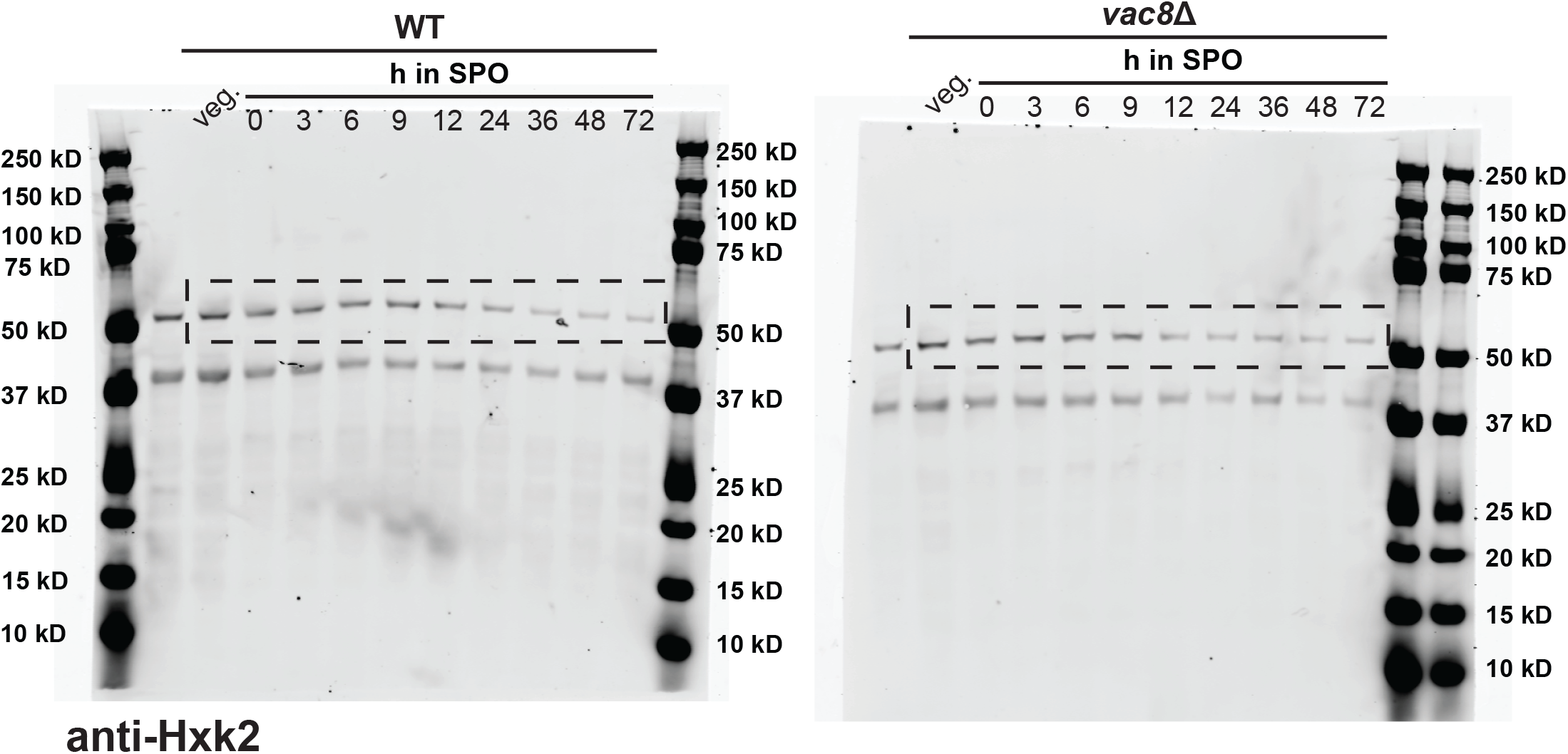
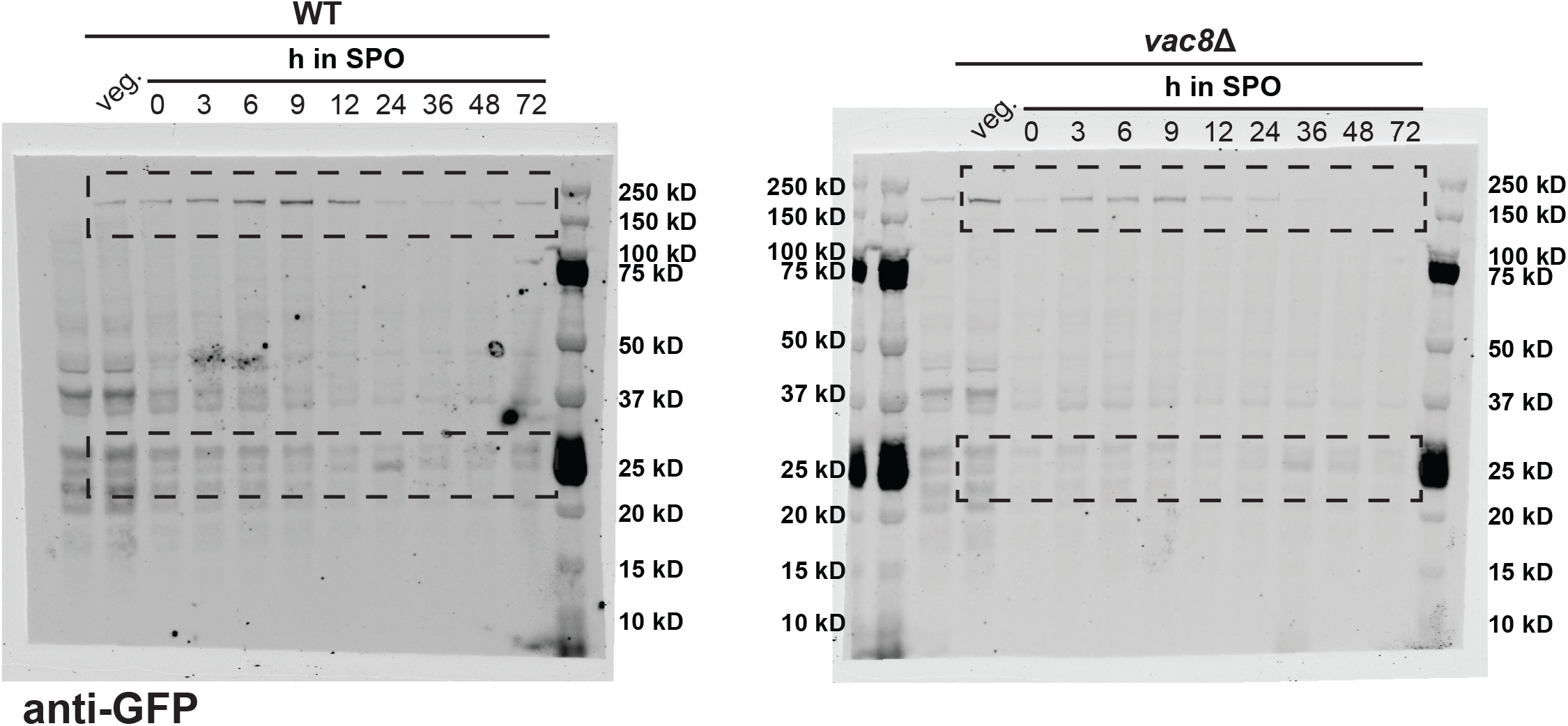
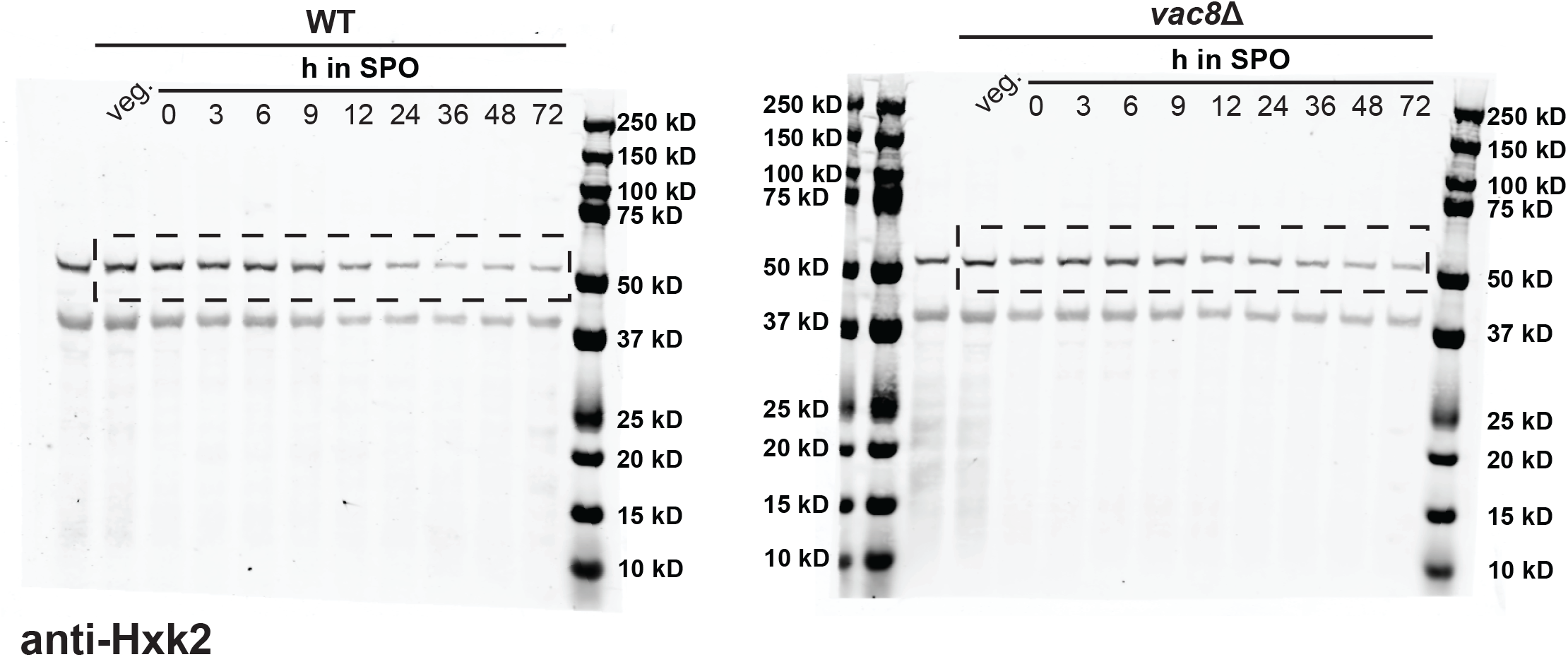

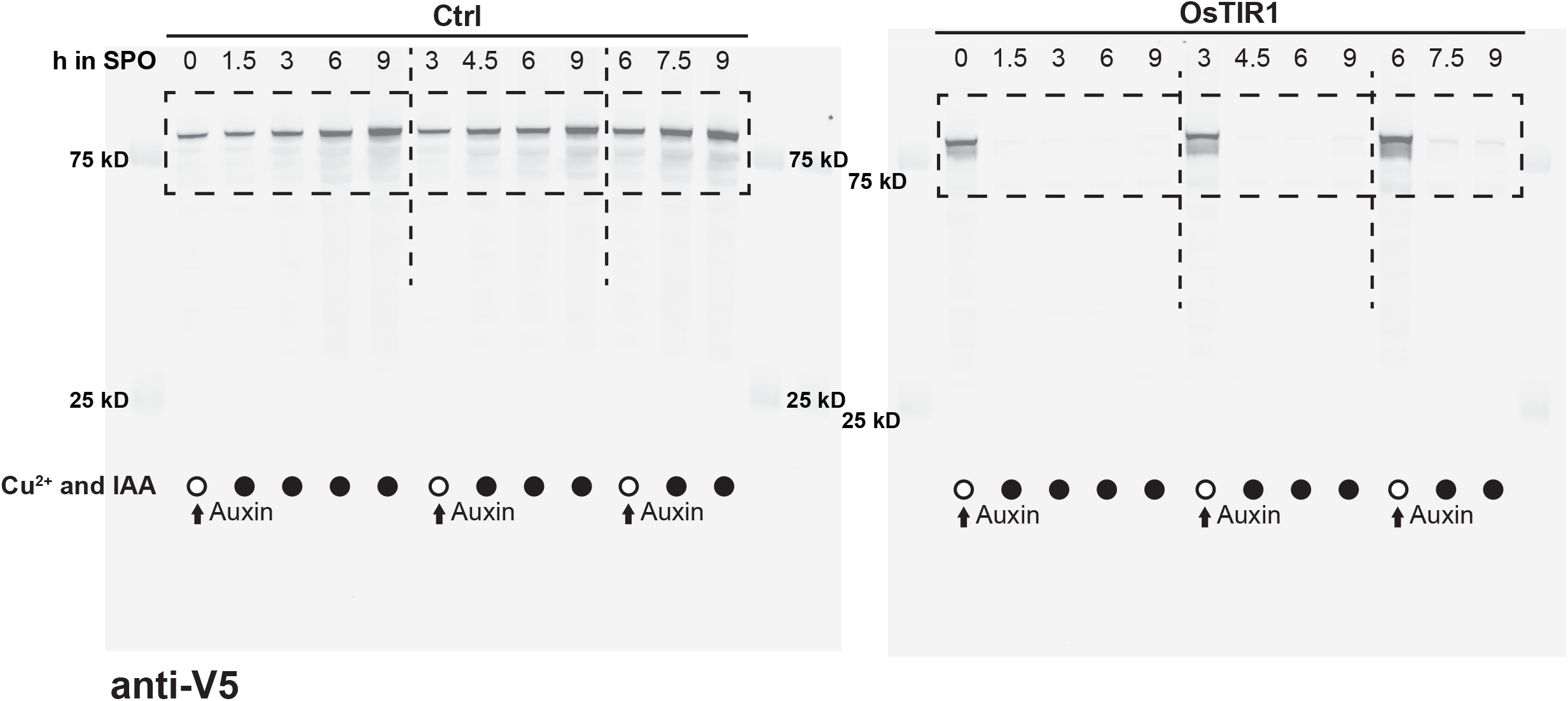
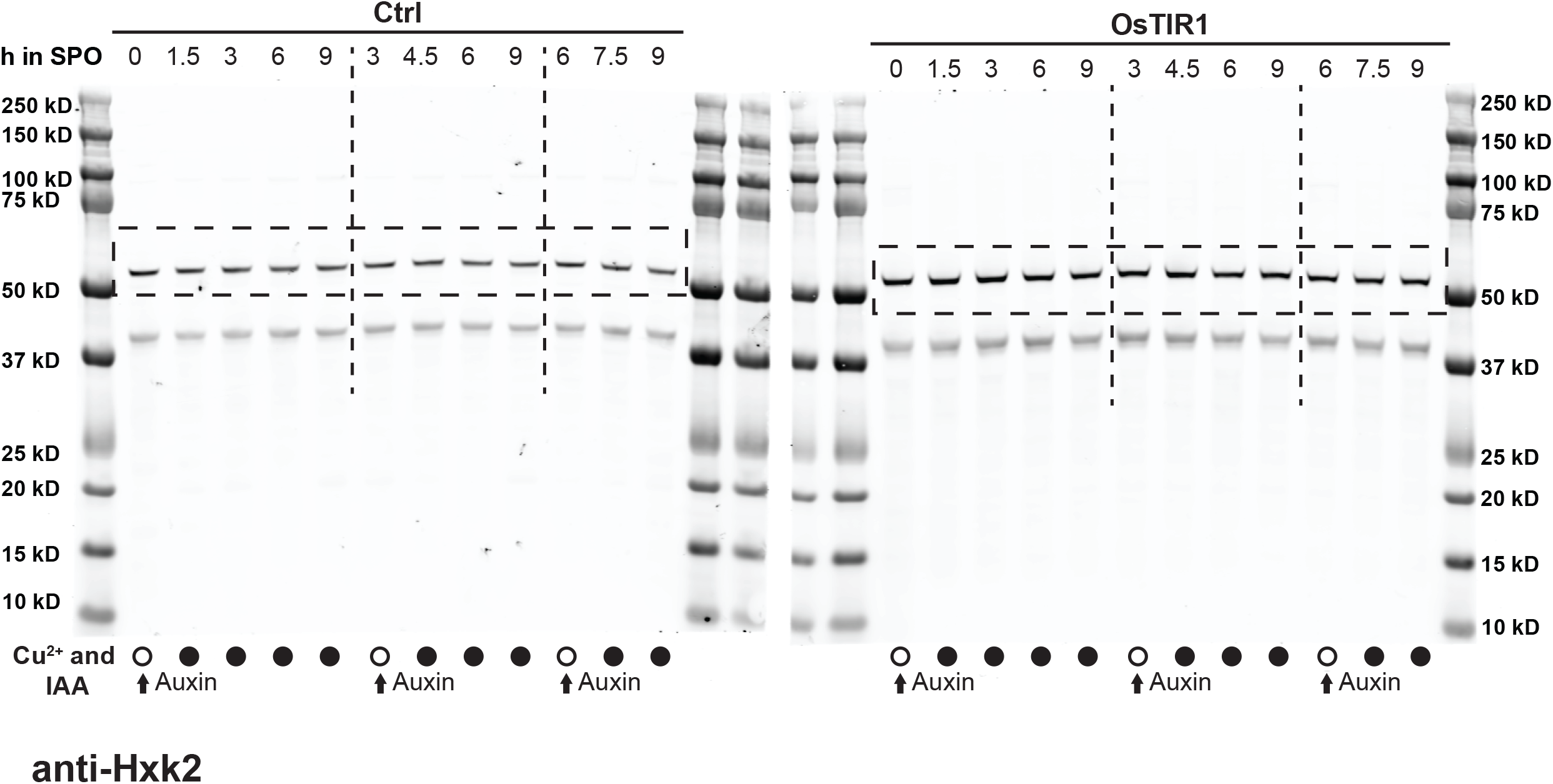

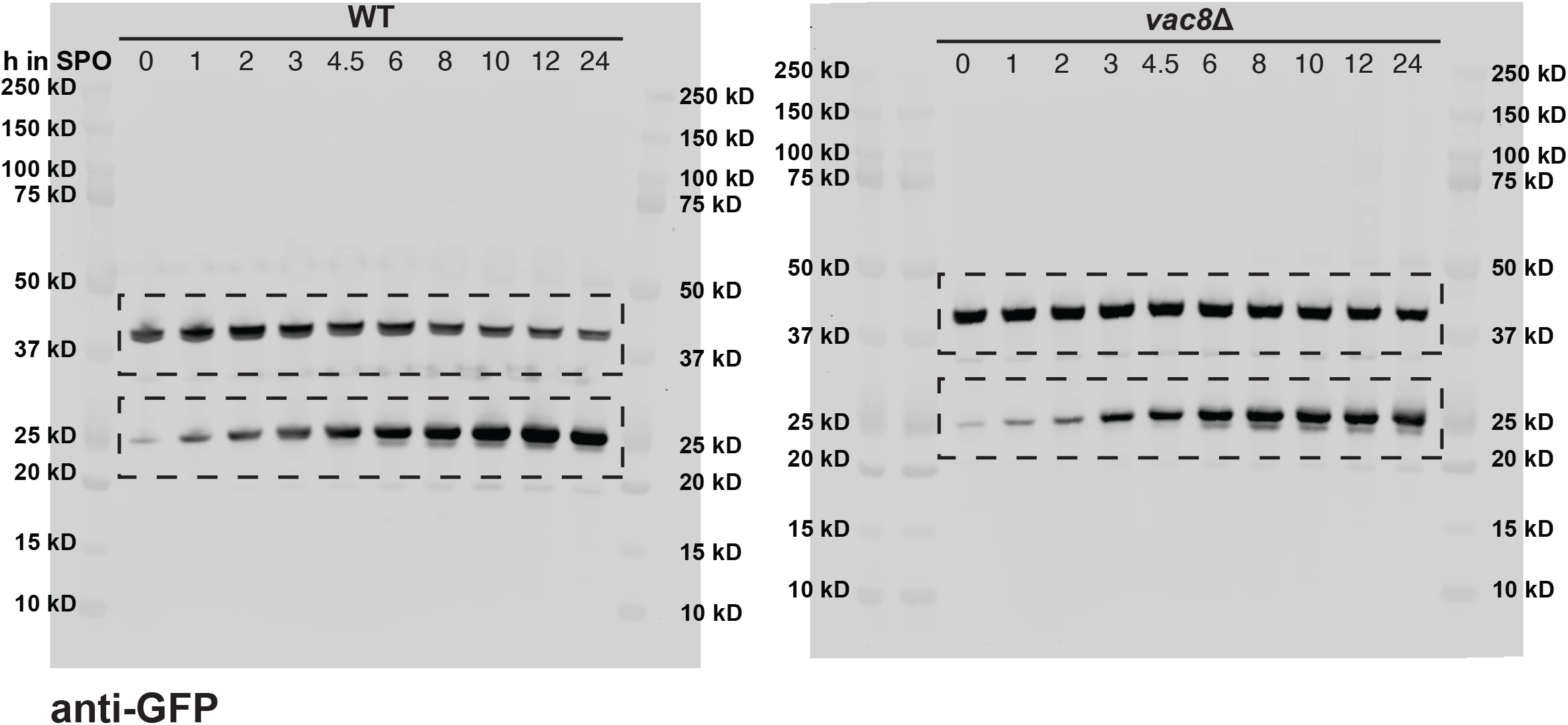
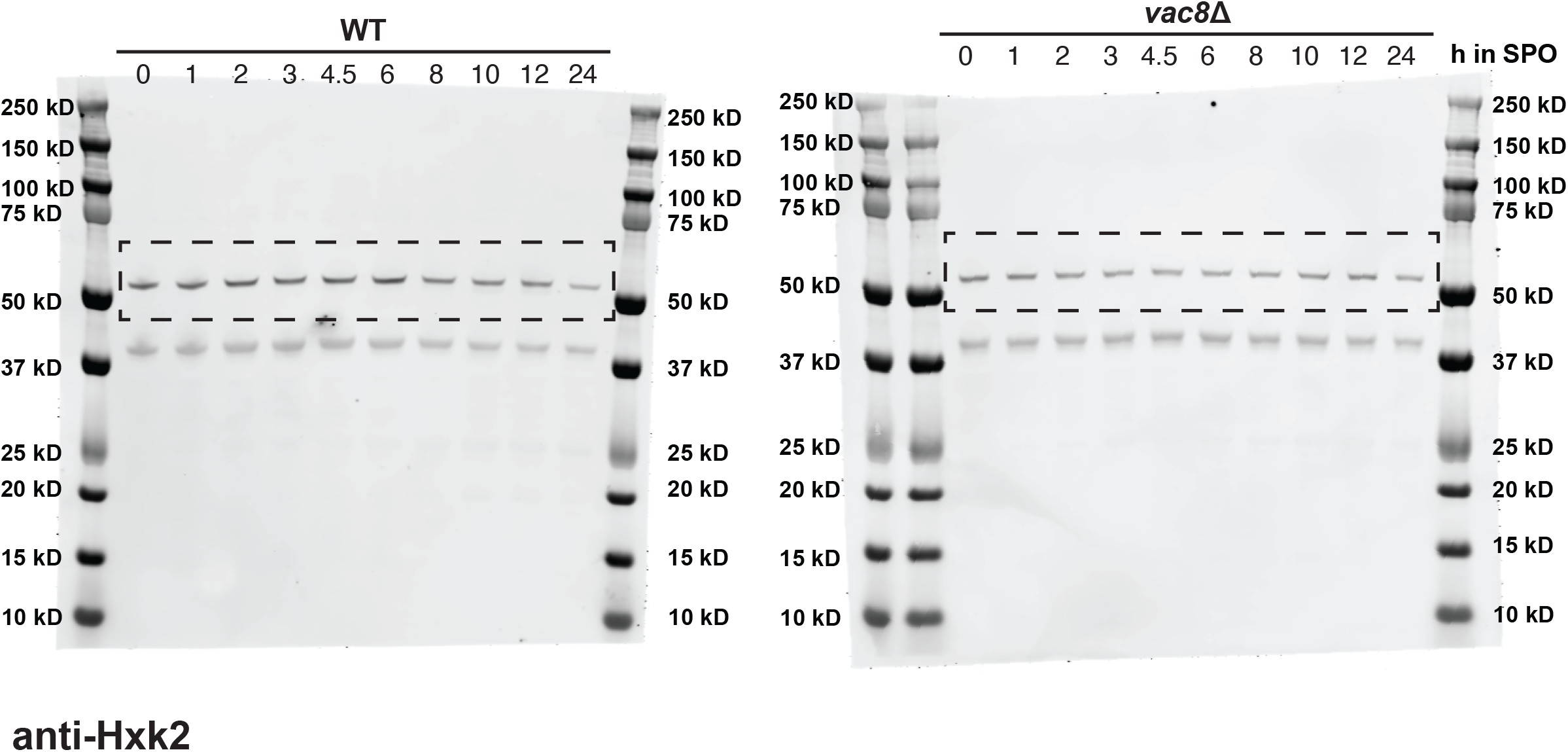

## Acknowledgments

We thank members of the Ünal and Brar laboratories (Department of Molecular and Cell Biology, University of California, Berkeley, CA, USA) for their support regarding this research, especially Cyrus Ruediger, Thomas Moss, Silvan Spiri, and Him Visitvoranat for their thoughtful comments on the manuscript. We thank members of the Charles Boone (Department of Molecular Genetics, Donnelly Centre, University of Toronto; RIKEN Center for Sustainable Resource Science) and Brenda Andrews laboratory (Department of Molecular Genetics, Donnelly Centre, University of Toronto) for their help on the genetic screen. We thank Grant Brown (Department of Molecular Genetics, Donnelly Centre, University of Toronto) for gifting us the “fixed” BY strains. Research was supported by funds from the National Institutes of Health (R01AG071801), John Templeton Foundation (63842), Astera Institute, and Shurl and Kay Curci Foundation (to E. Ünal); NIH training grant (T32 GM007232) and a National Science Foundation Graduate Research Fellowship (DGE 2146752) (to B.S. Styler).

## Author contributions

conceptualization, T. Xiao and E. Ünal; methodology and formal analysis, T. Xiao and B.S. Styler; investigation, T. Xiao and B.S. Styler; visualization, T. Xiao and E. Ünal; funding acquisition, B.S. Styler and E. Ünal; supervision, E. Ünal; project administration, E. Ünal; writing—original draft, T. Xiao and E. Ünal; writing—review and editing, T. Xiao, and E. Ünal.

---

**Figure S1.**
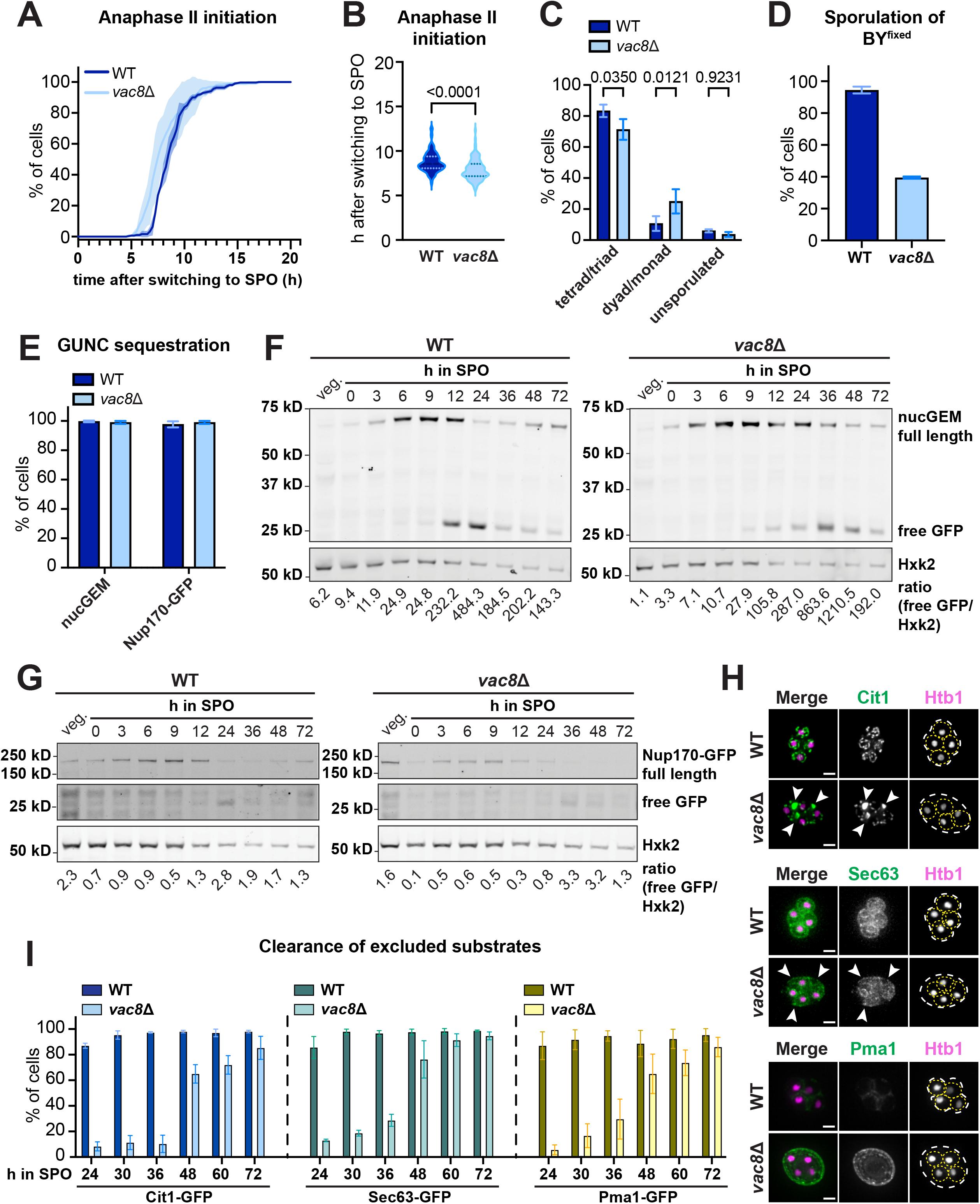
*VAC8* deletion does not inhibit meiotic progression or GUNC sequestration, but perturbs mega-autophagy (related to Fig. 1). **(A)** The percentage of WT and *vac8*Δ cells that had initiated anaphase II over time after switching to SPO, quantified from time-lapse images. Anaphase II initiation was determined by the formation of four dividing gamete nuclei, visualized by Htb1-mCh. Shaded region indicates SD of three replicates, n = 100 cells per replicate. (B) Violin plot showing the time it took for WT and *vac8*Δ cells to enter anaphase II after switching to SPO. n = 300 cells. (C) The percentage of tetrads/triads, dyads/monads, and unsporulated cells formed by WT and *vac8*Δ cells after 24 h in SPO. Error bars = SD of three replicates, n = 100 cells per replicate. (D) The percentage of WT and *vac8*Δ cells of “fixed” BY background that had sporulated at 24 h in SPO. Error bars = SD of two replicates. n = 100 cells per replicate. (E) Quantification of Fig. 1B and Fig. 1D showing the percentage of WT and *vac8*Δ cells that sequestered indicated substrates in the GUNC during anaphase II. Error bars = SD of three replicates. n = 100 cells per replicate. (F and G) Western blot analysis of GFP-tagged nucGEM (F) and Nup170 (G) in WT and *vac8*Δ cells undergoing vegetative growth (veg.) or progressing through gametogenesis. Samples were collected at indicated time points in SPO. Hxk2 was used as a loading control. The ratio of free GFP to Hxk2 band intensity in each sample is indicated at the bottom. **(H)** Widefield microscopy images of sporulated WT and *vac8*Δ cells at 24 h in SPO. Chromatin was visualized with Htb1-mCherry. Mitochondria, ER, and the plasma membrane were visualized by GFP-labelled Cit1, Sec63, and Pma1, respectively. Scale bar = 2 µm. The asci are outlined in white, and spores are outlined in yellow. White arrowheads indicate uninherited organelle remnants in *vac8*Δ mutants. Images are shown as MIPs of z-stacks. Scale bar = 2 µm. **(I)** Quantification of (G) showing the percentage of sporulated WT and *vac8*Δ cells that had cleared indicated uninherited organelle contents at indicated time points in SPO. Error bars = SD of three replicates. n = 100 cells per replicate. For statistical analysis information related to (B) and (C), see Table S5. The p-values are indicated in the graph.

**Figure S2.**
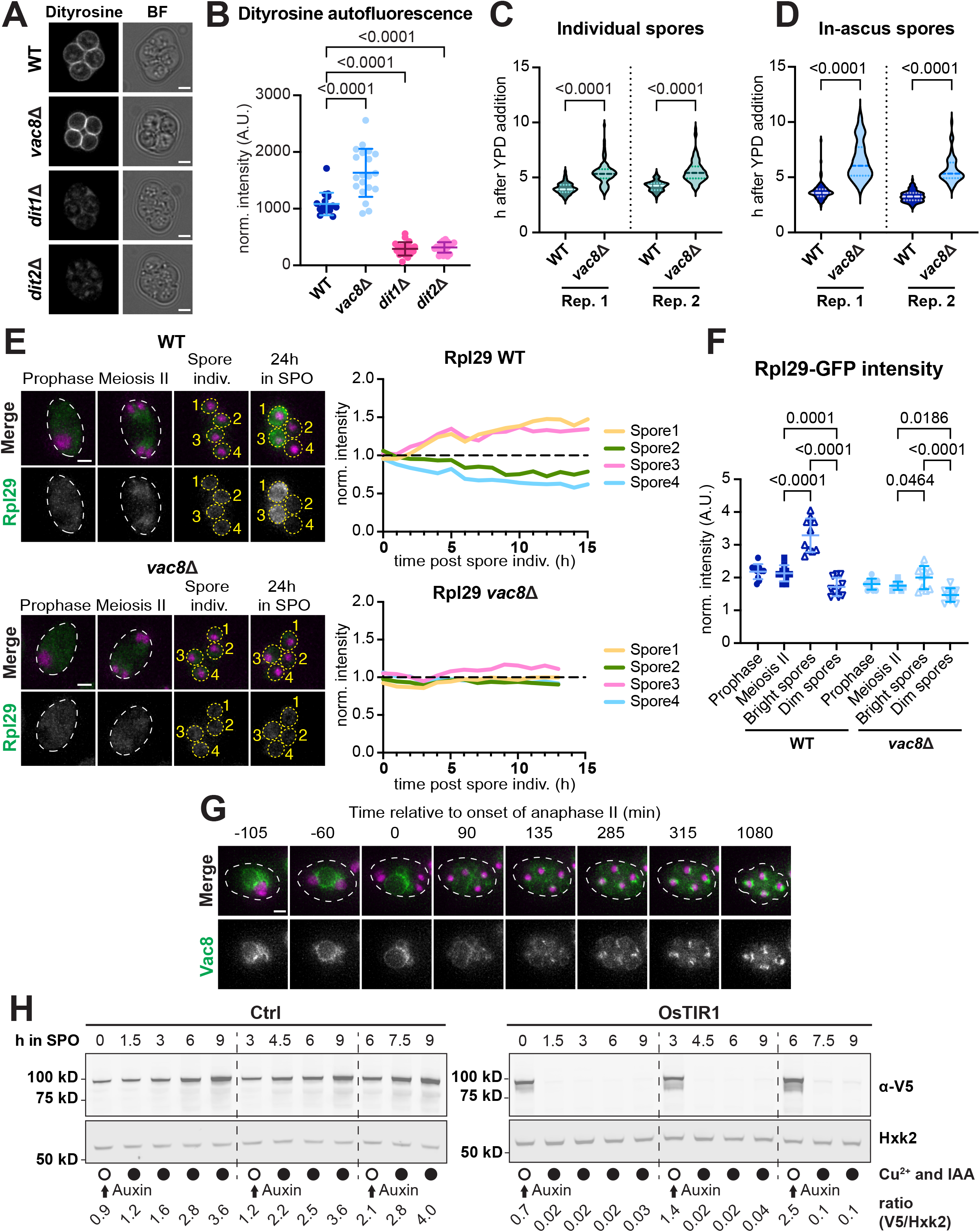
The deletion of *VAC8* affects spore competence, and Vac8 can be depleted during gametogenesis. (related to Fig. 2 and 3). **(A)** Widefield microscopy images of dityrosine autofluorescence in sporulated wild type and indicated mutants. Sporulated asci were collected at 24 h in SPO, resuspended 5% NH_4_OH for alkalization. Image represents a single z-slice. BF, brightfield. Scale bar = 2 µm. (**B**) Quantification of (A) showing the normalized intensity of dityrosine autofluorescence in WT and indicated mutants. Each dot indicates an individual sporulated ascus. n = 20 asci. (**C and D**) Quantification of Fig. 2F showing the time it took for free (C) or in-ascus spores (D) to germinate after switching to YPD. n = the number of spores that had germinated in Fig. 2 G and H. **(E)** Montage of WT and *vac8*Δ cells expressing endogenously labelled Rpl29-GFP at indicated stages during gametogenesis. Chromatin was visualized with Htb1-mCherry (shown in magenta in merged images). The precursor cells or asci are outlined in white, and spores are outlined in yellow. Images are shown as MIPs of z-stacks. Scale bar = 2 µm. The line plots on the right depict normalized Rpl29 fluorescence intensity in individual spores at each time point after spore individualization. Each line represents the respective spore labelled in the microscopy image. The time point of spore individualization was defined as 0 h. Fluorescence intensity was normalized to that at 0 h, and the dashed black line indicates normalized intensity = 1.0. **(F)** Quantification of the normalized Rpl29 fluorescence intensity in WT or *vac8*Δ cells during prophase, anaphase II, and in bright versus dim spores after 24 h in SPO. Each symbol represents an individual cell or spore, n = 10 sporulated cells. Error bars = SD across all individual cells or spores within each group. **(G)** Montage of cells expressing endogenously tagged Vac8-GFP progressing through gametogenesis. Chromatin was visualized with Htb1-mCherry. The time point depicting anaphase II onset was defined as 0 min. Images are shown as MIPs of z-stacks. Scale bar = 2 µm. **(H)** Western blot analysis of Vac8-3V5 in Ctrl or OsTIR1 cells progressing through gametogenesis. Samples were collected at indicated time points in SPO. Cu^2+^ and 5-Ph-IAA were added to the sporulating culture at indicated time points after switching to SPO. Hxk2 was used as a loading control. The ratio of V5 to Hxk2 band intensity in each sample is indicated at the bottom. For statistical analysis information related to (B), (C), (D), and (F), see Table S5. The p-values are indicated in the graph.

**Figure S3.**
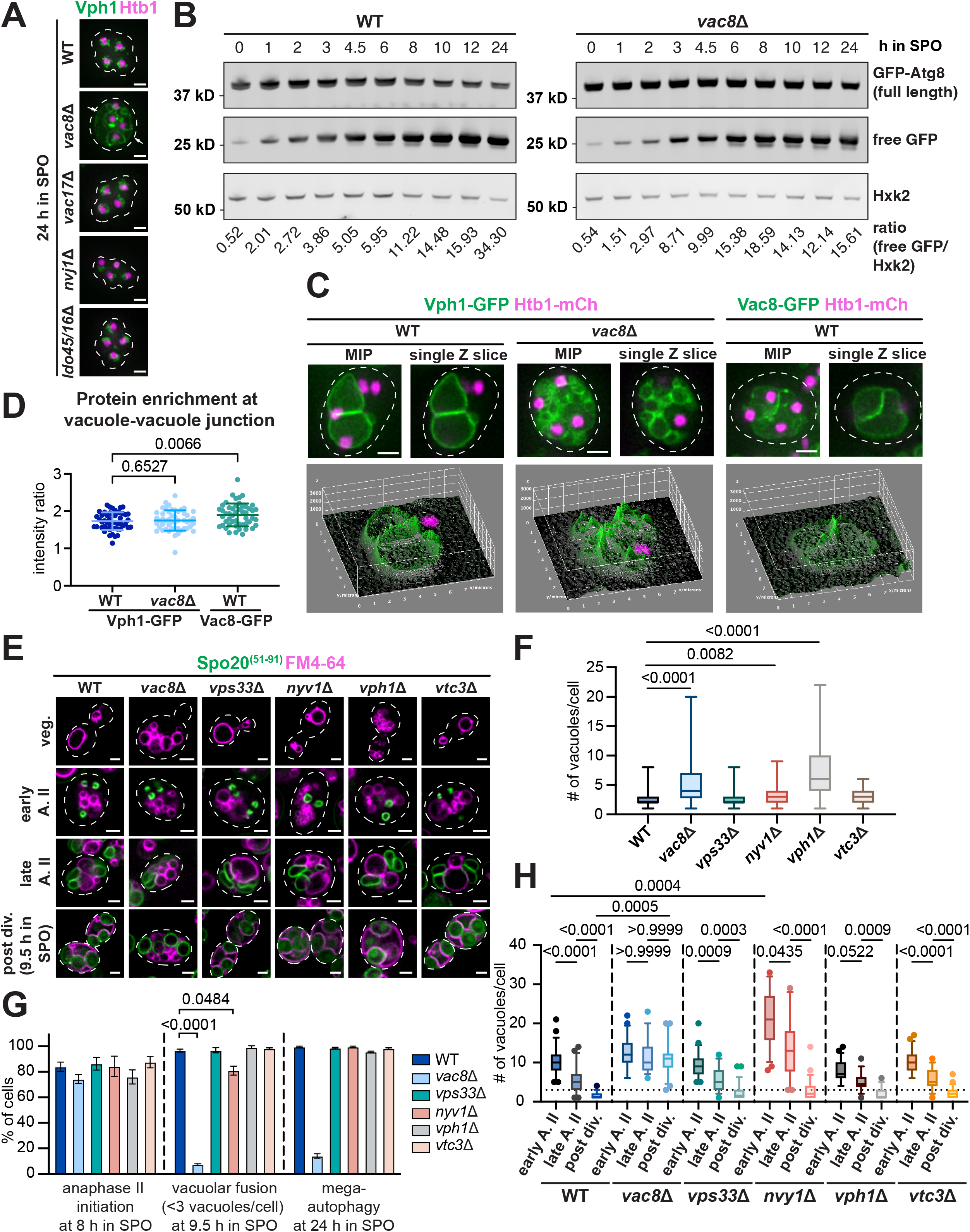
Several known Vac8-associated proteins were dispensable for meiotic vacuolar remodeling (related to Fig. 5). **(A)** Widefield microscopy images of sporulated WT and indicated mutants at 24 h in SPO. Vacuoles were visualized with Vph1-GFP, and chromatin was visualized with Htb1-mCherry. White arrow indicates the unlysed precursor vacuoles. Images are shown as MIPs of z-stacks. Scale bar = 2 µm. **(B)** Western blot analysis of GFP-tagged Atg8 in WT and *vac8*Δ cells progressing through gametogenesis. Samples were harvested at indicated time points in SPO. Hxk2 was used as a loading control. The ratio of free GFP to Hxk2 band intensity in each sample is indicated at the bottom. **(C)** Widefield microscopy images showing the vacuole-vacuole contacts in WT and *vac8*Δ cells during late meiosis II. Vph1 or Vac8 was labelled with GFP, and chromatins were visualized with Htb1-mCherry. Images are shown as MIPs of z-stacks or single z-slices. Scale bar = 2 µm. The 3D surface plot represents GFP fluorescence intensity across the entire single z-plane. **(D)** Quantification of (C) showing the GFP intensity of indicated proteins at vacuole-vacuole contacts normalized to non-contacting vacuolar membrane. Each dot indicates an individual vacuole. n = 50 vacuoles. **(E)** Widefield microscopy images of wild type and indicated mutants undergo vegetative growth (veg.) or progressing through gametogenesis at indicated stages. Precursor vacuoles were visualized by FM4-64 staining prior to switching to SPO. The PSM was marked by GFP-SPO20^(51-91)^. The stages at anaphase II were determined by PSM morphology. A. II, anaphase II. Post-division (post div.) samples were harvested at 9.5 h in SPO. Image represents a single z-slice. Scale bar = 2 µm. **(F)** Quantification of (E) showing the number of vacuoles per cell in WT and indicated mutants that undergo vegetative growth. Error bars indicate the range (min to max) of values. **(G)** The percentage of WT and indicated mutants that had initiated anaphase II at 8 h in SPO, completed vacuolar fusion (defined by ≤ 3 vacuoles per cell), and completed mega-autophagy at 24 h in SPO. Error bars = SD of three replicates, n = 100 cells per replicate. **(H)** Quantification of (E) showing the number of vacuoles per cell in WT and indicated mutants progressing through gametogenesis at indicated stages. Error bars indicate the 95% confidence interval, and outliers are shown as dots. The dotted black line indicates vacuole number = 3 per cell. For statistical analysis information related to (D), (F), (G), and (H), see Table S5. For (D) and (H), the p-values are indicated in the graph. For (F) and (G), the p-values are indicated for comparisons with p ≤ 0.05. For cross-genotype comparisons with WT in (H), only comparisons with p ≤ 0.05 are indicated.

**Table S1.**
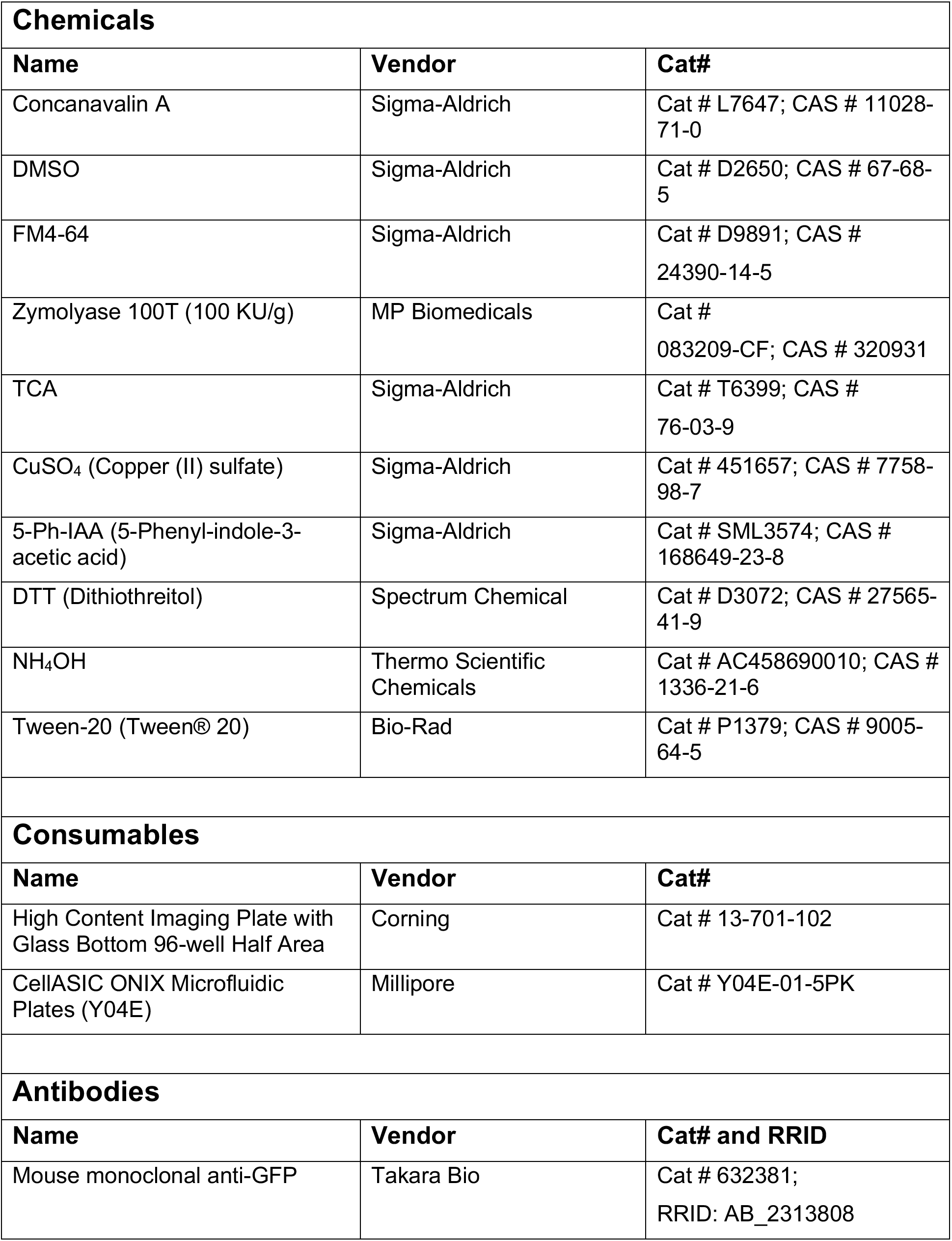
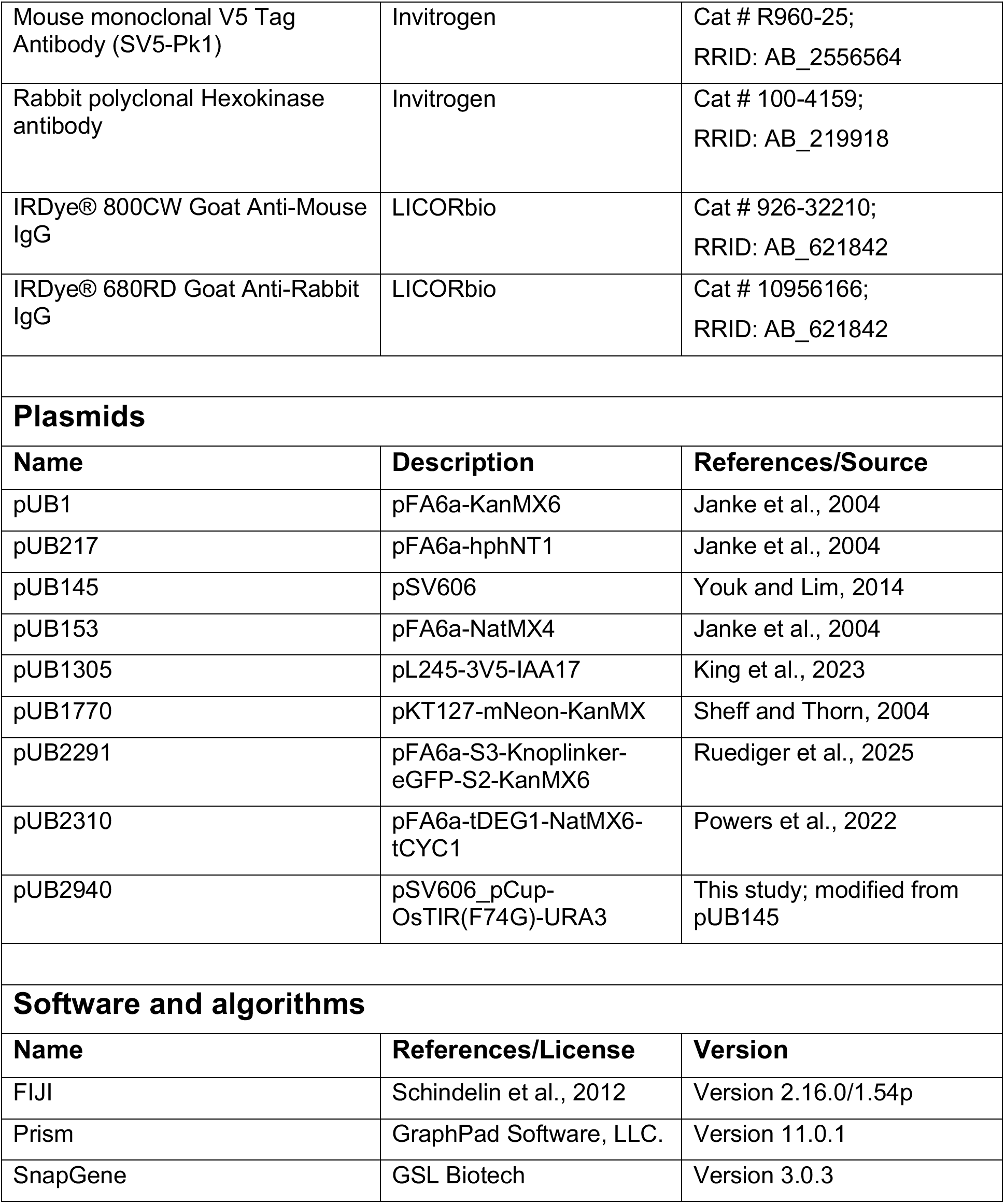
Chemicals, antibodies, plasmids, and software used in this study.

**Table S2.**
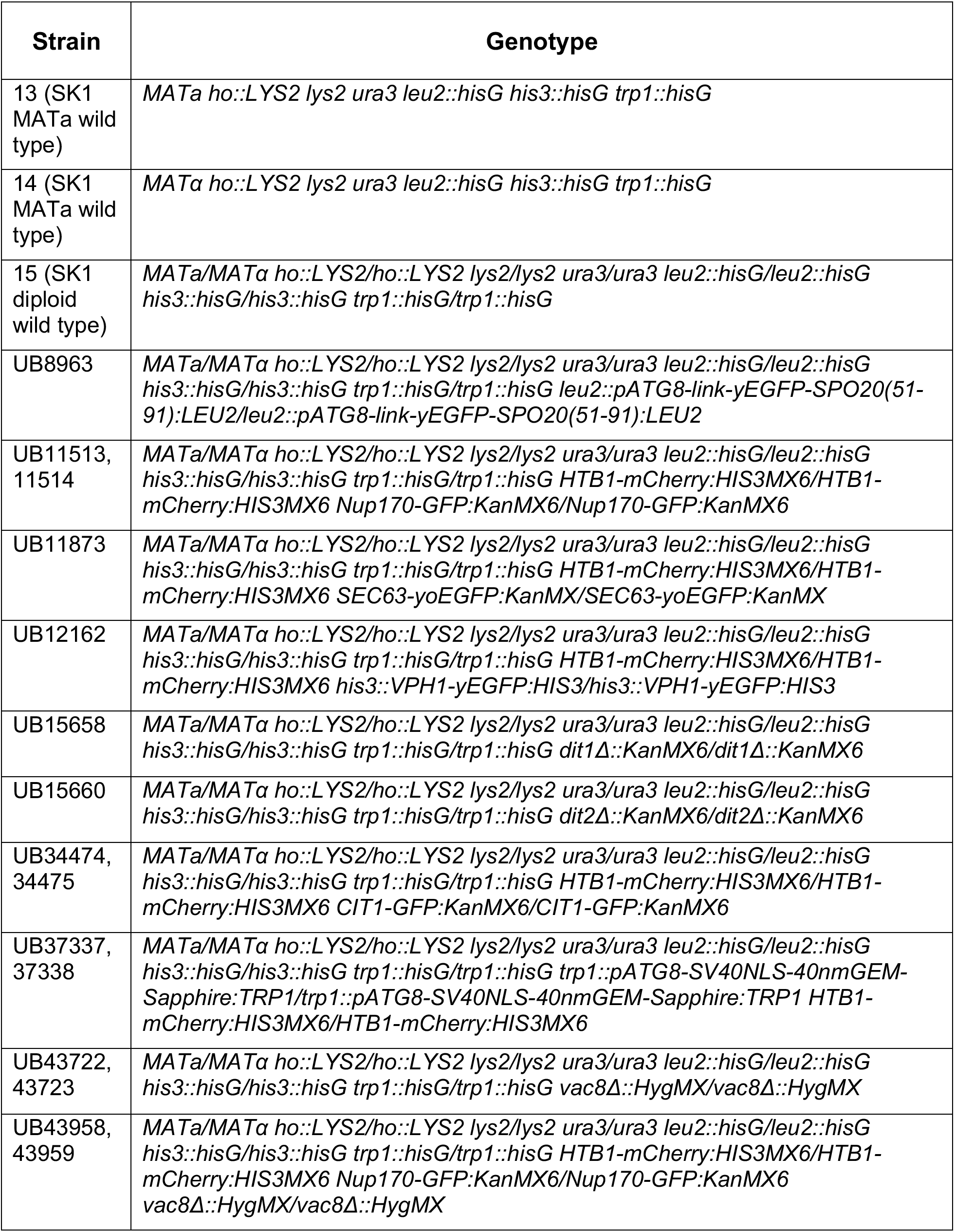
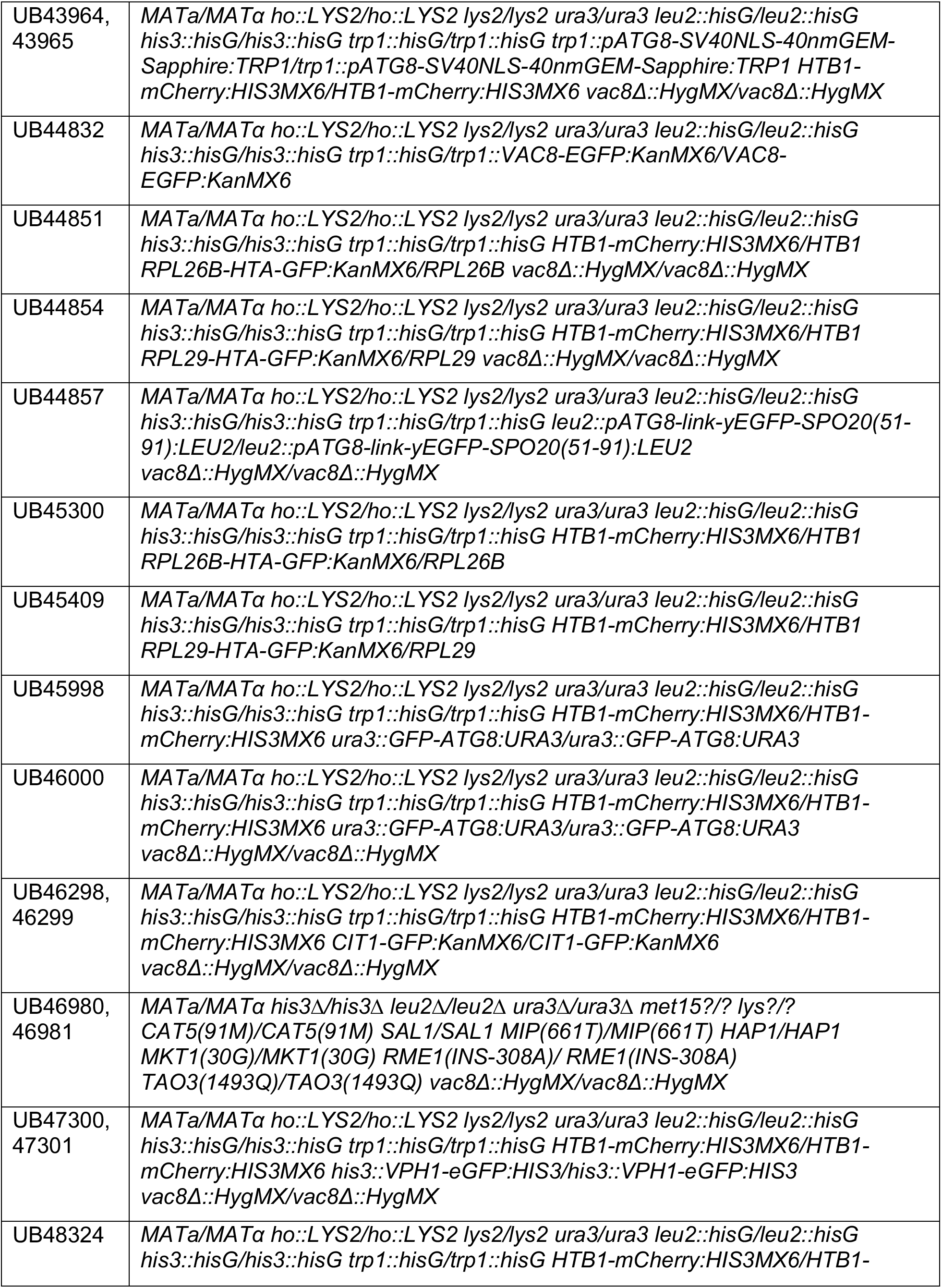
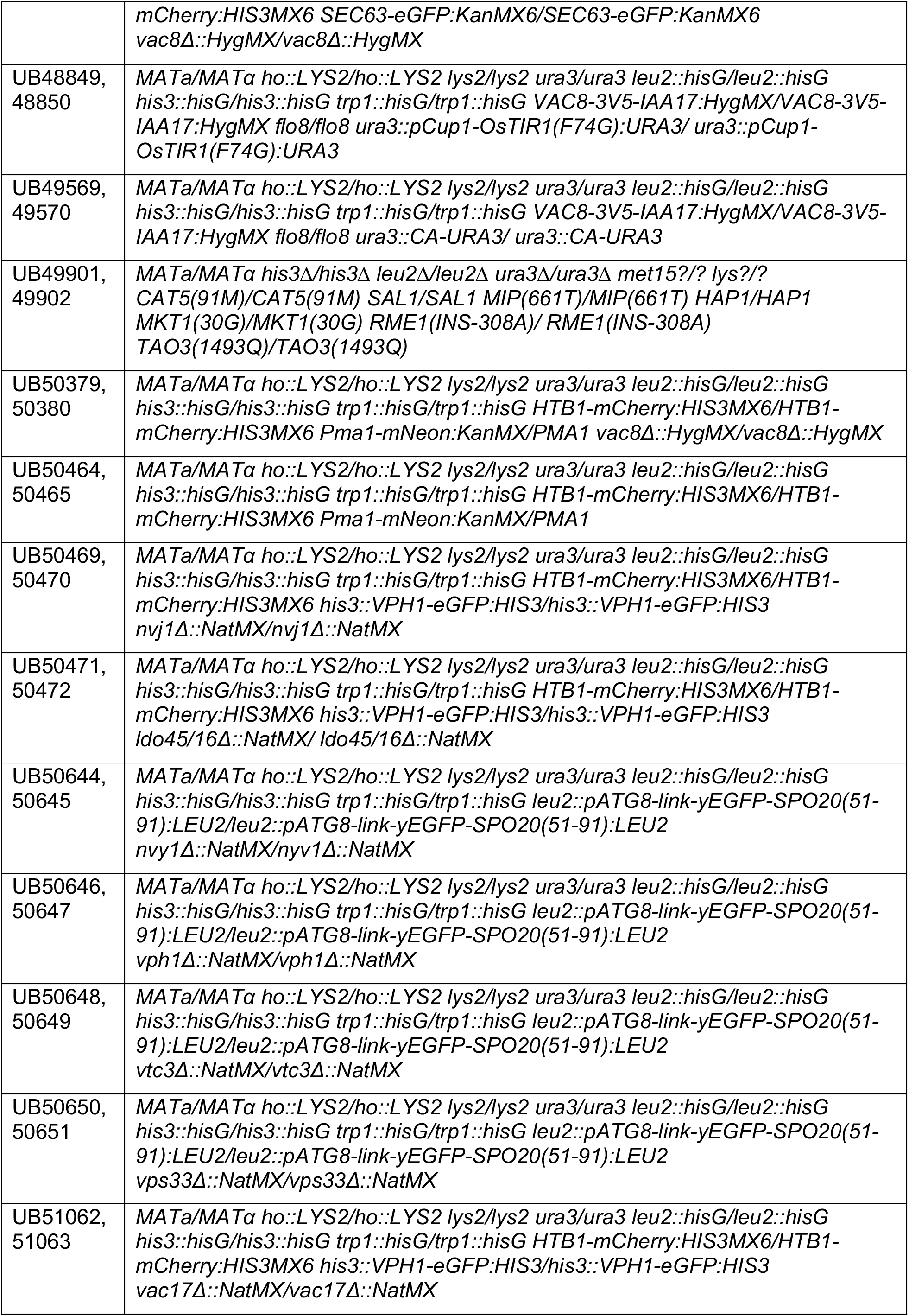
Yeast strains used in this study.

**Table S3.**
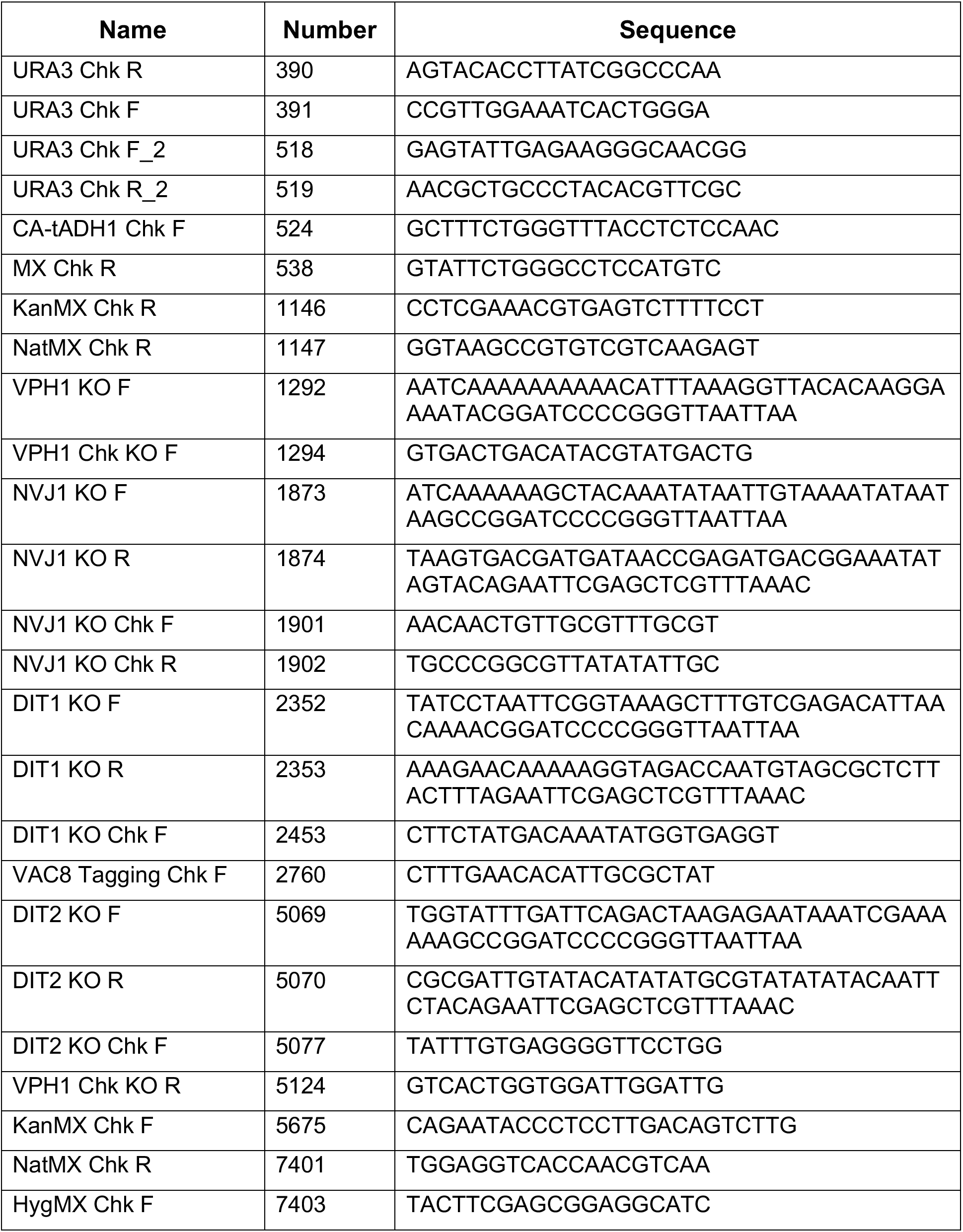
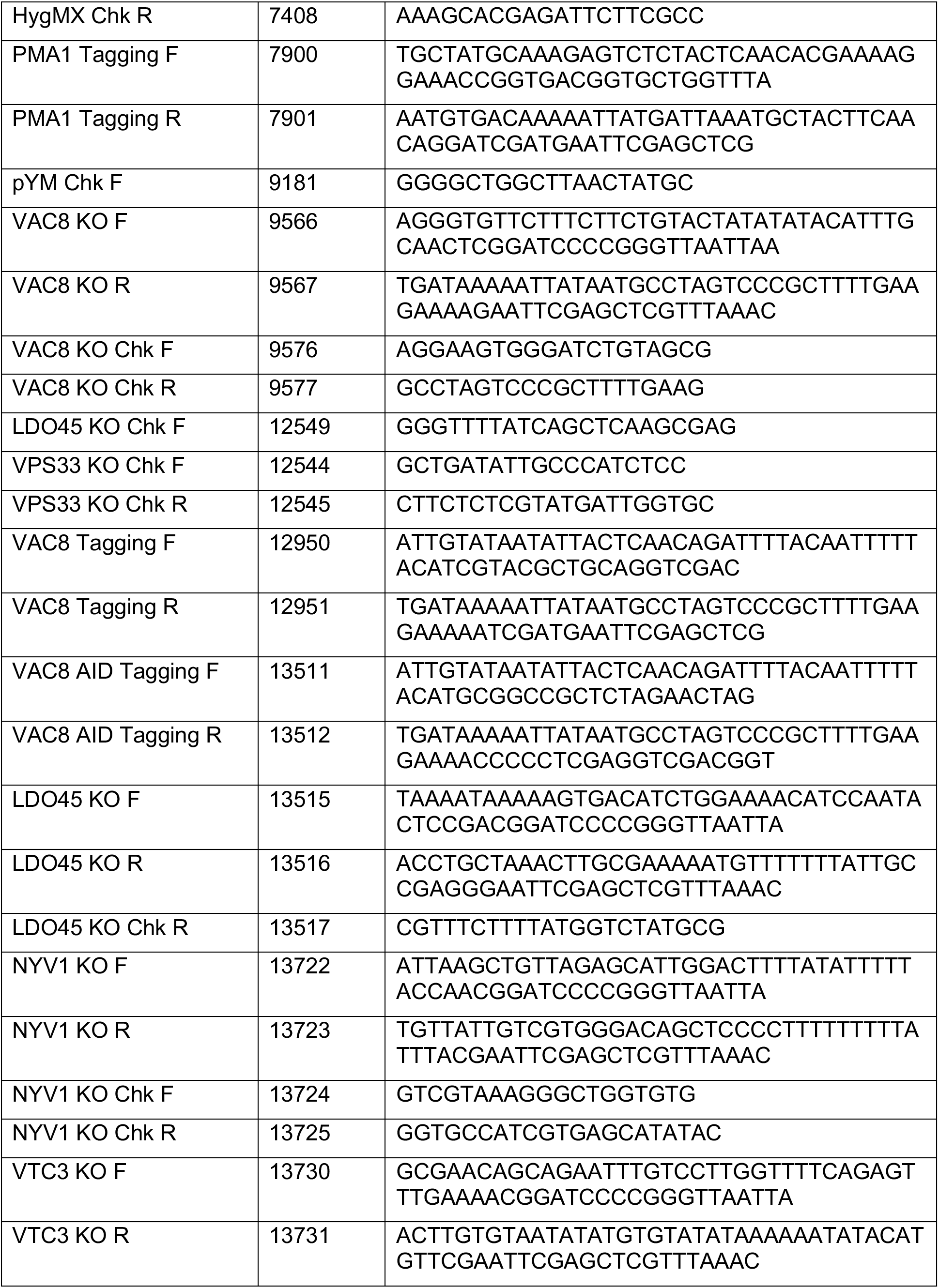
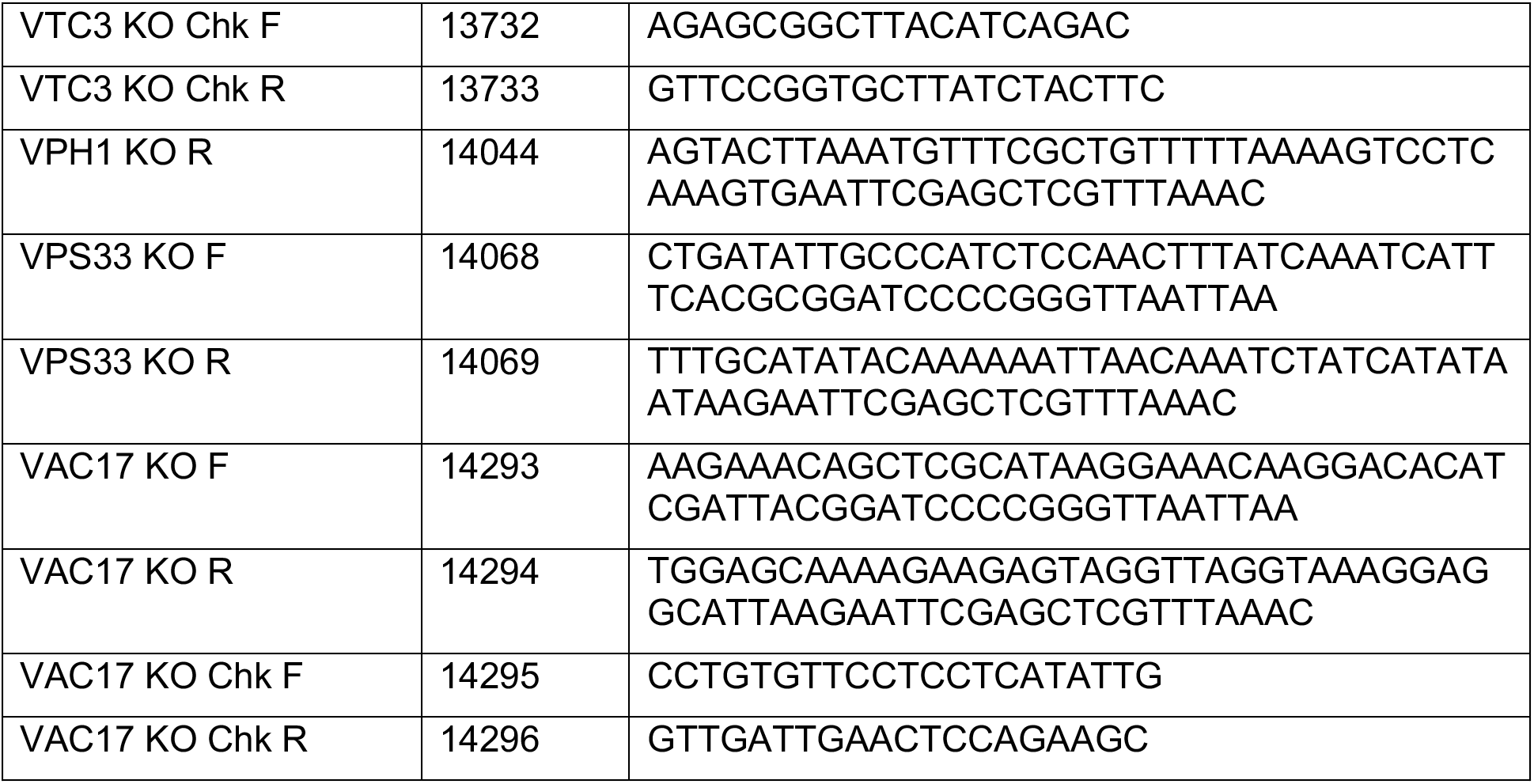
Oligos used in this study.

**Table S4.** Imaging Conditions.

| **Figure** | **Experiment type** | **Exposures** | **Sectioning** | **Acquisition frequency and duration** | **Image projection** |
| --- | --- | --- | --- | --- | --- |
| 1B, 1E, 2I, and S2E | Time-lapse, 96-well plate | mCherry: 10%T, 25 ms;<br>FITC: 10%T, 25 ms;<br>POL: 32% T, 100 ms. | 1 μm, 8 sections;<br>60x obj | 15 min/frame, 20 h in total | Max-intensity projection |
| 2A and 3E | Still images, live cell, slides | TRITC: 100%T, 25 ms;<br>POL: 32% T, 100 ms. | 1 μm, 8 sections;<br>60x obj | N/A | Max-intensity projection |
| 2F and 4D | Time-lapse, 96-well plate | POL: 32% T, 100 ms. | 1 μm, 8 sections;<br>60x obj | 5 min/frame, 10 h in total | Single z-slice |
| 5C and S3E | Still images, live cell, slides | TRITC: 100%T, 25 ms;<br>FITC: 100%T, 25 ms;<br>POL: 32% T, 100 ms. | 1 μm, 8 sections;<br>60x obj | N/A | Single z-slice |
| 5F | Time-lapse, Cell ASIC microfluidics platform | mCherry: 10%T, 25 ms;<br>FITC: 10%T, 25 ms;<br>POL: 32% T, 100 ms. | 1 μm, 8 sections;<br>60x obj | 15 min/frame, 20 h in total | Max-intensity projection |
| S1H and S3A | Still images, live cell, slides | mCherry: 100%T, 25 ms;<br>FITC: 100%T, 25 ms;<br>POL: 32% T, 100 ms. | 1 μm, 8 sections;<br>60x obj | N/A | Max-intensity projection |
| S2A | Still images, live cell, slides | FITC: 100%T, 25 ms;<br>POL: 32% T, 100 ms. | 1 μm, 8 sections;<br>60x obj | N/A | Max-intensity projection |
| S2G | Time-lapse, 96-well plate | mCherry: 10%T, 25 ms;<br>FITC: 32%T, 25 ms;<br>POL: 32% T, 100 ms. | 1 μm, 8 sections;<br>60x obj | 15 min/frame, 20 h in total | Max-intensity projection |
| S3C | Still images, live cell, slides | mCherry: 100%T, 25 ms;<br>FITC: 100%T, 25 ms;<br>POL: 32% T, 100 ms. | 1 μm, 8 sections;<br>60x obj | N/A | As indicated |

**Table S5.** Quantification and statistical analysis information.

| Figure | Replicate # | N per replicate | Statistical analysis method | Error bar |
| --- | --- | --- | --- | --- |
| 1C, 1D, 1F, 1G, 2B, 3C, 3D, 5B, 5G, S1A, S1E, and S1I | 3 | 100 cells | N/A | SD |
| 2D, 2E, 4B, and 4C | 3 | 2D Rep. 1<br>WT: 91 colonies;<br>*vac8Δ*: 91 colonies | Welch's ANOVA with Dunnett's T3 multiple comparisons test (not assuming no equal SD across groups) | SD |
|  |  | 2D Rep. 2<br>WT: 91 colonies;<br>*vac8Δ*: 86 colonies |  |  |
|  |  | 2D Rep. 3<br>WT: 96 colonies;<br>*vac8Δ*: 90 colonies |  |  |
|  |  | 2E Rep. 1<br>WT: 94 colonies;<br>*vac8Δ*: 88 colonies |  |  |
|  |  | 2E Rep. 2<br>WT: 96 colonies;<br>*vac8Δ*: 82 colonies |  |  |
|  |  | 2E Rep. 3<br>WT: 96 colonies;<br>*vac8Δ*: 85 colonies |  |  |
|  |  | 4B Rep. 1<br>WT: 90 colonies;<br>*vac8Δ*: 90 colonies |  |  |
|  |  | 4B Rep. 2<br>WT: 89 colonies;<br>*vac8Δ*: 94 colonies |  |  |
|  |  | 4B Rep. 3<br>WT: 79 colonies;<br>*vac8Δ*: 83 colonies |  |  |
|  |  | 4B Rep. 1<br>WT: 96 colonies;<br>*vac8Δ*: 94 colonies |  |  |
|  |  | 4B Rep. 2<br>WT: 95 colonies;<br>*vac8Δ*: 91 colonies |  |  |
|  |  | 4B Rep. 3<br>WT: 94 colonies;<br>*vac8Δ*: 93 colonies |  |  |
| 2G, 2H, 4E, 4F | 2 | 100 cells | N/A | SD |
| 2J and S2F | N/A | 10 asci | Two-way repeated-measures ANOVA (mixed-effects model), with genotype and time as factors, and Geisser-Greenhouse correction applied. Holm-Šidák's multiple comparisons test was used to compare genotypes at each time point. | SD |
| 4G, 4H, S2C, and S2D | 2 | 4G Rep. 1<br>WT: 100 spores;<br>*vac8Δ*: 100 spores | Welch's ANOVA with Dunnett's T3 multiple comparisons test | N/A |
|  |  | 4G Rep. 2<br>WT: 100 spores;<br>*vac8Δ*: 100 spores |  |  |
|  |  | S2C Rep. 1<br>WT: 100 spores;<br>*vac8Δ*: 100 spores |  |  |
|  |  | S2C Rep. 2<br>WT: 100 spores;<br>*vac8Δ*: 99 spores |  |  |
| 4H, S2D |  | 4H Rep. 1<br>WT: 100 asci;<br>*vac8Δ*: 84 asci | Kruskal-Wallis test with Dunn's multiple comparisons test (as the data did not follow a normal distribution) | N/A |
|  |  | 4H Rep. 2<br>WT: 100 asci;<br>*vac8Δ*: 84 asci |  |  |
|  |  | S2D Rep. 1<br>WT: 100 asci;<br>*vac8Δ*: 58 asci |  |  |
|  |  | S2D Rep. 2<br>WT: 100 spores,<br>*vac8Δ*: 40 spores |  |  |
| 5D | N/A | 50 cells | Kruskal-Wallis test with Dunn's multiple comparisons test | SD |
| 5E | N/A | All vacuoles from 50 cells | Kruskal-Wallis test with Dunn's multiple comparisons test | min to max |
| S1B | 3 | 100 cells | Unpaired t-test | N/A |
| S1C | 3 | 100 cells | Two-way repeated-measures ANOVA (mixed-effects model), with genotype and time as factors, and Geisser-Greenhouse correction applied. Šídák's multiple comparisons test was used to compare genotypes for each phenotype. | SD |
| S1D | 2 | 100 cells | N/A | SD |
| S2B | 3 | 20 asci | Welch's ANOVA with Dunnett's T3 multiple comparisons test | SD |
| S3D | N/A | 50 vacuoles | Kruskal-Wallis test with Dunn's multiple comparisons test. | SD |
| S3F | N/A | 150 cells | Kruskal-Wallis test with Dunn's multiple comparisons test. | min to max |
| S3G | 3 | 100 cells | Welch's ANOVA with Dunnett's T3 multiple comparisons test | SD |
| S3H | N/A | 50 cells | Kruskal-Wallis test with Dunn's multiple comparisons test. | 95% confidence interval; outliers are shown as dots. |

